# Lipid utilization in skeletal muscle cells is modulated *in vitro* and *in vivo* by specific miRNAs

**DOI:** 10.1101/288035

**Authors:** Francesco Chemello, Francesca Grespi, Alessandra Zulian, Pasqua Cancellara, Etienne Hebert-Chatelain, Paolo Martini, Camilla Bean, Enrico Alessio, Ruggero Ferrazza, Paolo Laveder, Graziano Guella, Carlo Reggiani, Chiara Romualdi, Paolo Bernardi, Luca Scorrano, Stefano Cagnin, Gerolamo Lanfranchi

## Abstract

Skeletal muscle is composed by different myofiber types that can preferentially use glycolysis or lipids for ATP production. How fuel preference is specified in these post-mitotic cells is unknown. Here we show that miRNAs are important players in defining the myofiber metabolic profile. mRNA and miRNA signatures of all myofiber types obtained at single cell level unveiled fiber-specific regulatory networks and identified two master miRNAs that coordinately control myofiber fuel preference and mitochondrial morphology. Our work provides a complete and integrated myofiber type-specific catalogue of genes and miRNAs expressed and establishes miR-27a-3p and miR-142-3p as key regulators of lipid utilization in skeletal muscle.

**HIGHLIGHTS:**

- Transcriptional networking in single cells distinguished myofibers based on glycolytic or oxidative metabolism, regulated by specific miRNAs
- miR-27a-3p and −142-3p influence mitochondrial morphology
- miR-27a-3p improves lipid utilization and increases glycogen storage both *in vitro* and *in vivo*
- miR-142-3p reduces lipid utilization both *in vitro* and *in vivo*

## INTRODUCTION

Half of the body mass of healthy individuals is constituted by skeletal muscle that supports body movements and is crucial in intermediate metabolism. These anatomical and physiological features are shared throughout the Vertebrate phylum, where muscle exerts a chief role in regulation of whole body metabolism. As a metabolic organ, skeletal muscle regulates systemic glucose homeostasis. When glucose is not consumed as fuel, it is accumulated as glycogen by muscle, and this storage may be almost five times more efficient as in the liver (Jensen et al., 2011). However, during physiological changes such as, for example, fasting, muscle can rapidly adapt to use fatty acids (FAs) and amino acids as oxidative substrates. Not surprisingly, impairment in muscle metabolic plasticity and insulin sensitivity is a hallmark of several metabolic diseases including type II diabetes and obesity (Sears and Perry, 2015).

The flexibility of skeletal muscle in the utilization of carbohydrate or lipid is associated with the expression of different panels of structural proteins that ultimately define different contractile properties (Schiaffino and Reggiani, 2011). The overall physiological diversity and plasticity is based on the repertoire and proportion of different types of myofibers that compose a particular muscle in a particular physiological condition. In small mammals, slow muscles such as *soleus* are mostly composed of slow-oxidative myofibers that express the slow type 1 myosin heavy chain (MyHC) and fast-oxidative myofibers that express type 2A or 2X MyHC. Type 1, 2A, and 2X mouse myofibers are considered oxidative myofibers and prevalently utilize lipids to meet most of their energy requirements aerobically. Instead, fast muscles, such as *extensor digitorum longus* (EDL), *gastrocnemius*, and the *tibialis anterior*, are mostly composed by fast-glycolytic myofibers that express a predominance of type 2B MyHC and utilize glucose to produce most of energy anaerobically. Mixed with these four major types of myofibers, minor intermediate hybrid fibers expressing different combinations of MyHC isoforms can be found in different muscles.

Metabolic adaptation can also occur through the modulation of mitochondrial dynamics (Liesa and Shirihai, 2013; Schrepfer and Scorrano, 2016). In skeletal muscle, mitochondrial dynamics is tailored to the different myofiber types: mitochondria are elongated due to increased fusion in oxidative myofibers compared to glycolytic myofibers (Mishra et al., 2015). This layer of regulation can contribute to explain how muscle, a terminally differentiated and post-mitotic tissue, can rapidly adapt its metabolism.

Muscle adaptive capacity to specific physiological stimuli should rely on complex, ordered changes in the pattern of fiber gene expression. Indeed, gene expression profiles of muscles with different metabolic traits are different (Campbell et al., 2001; Wu et al., 2003) and sensor molecules exist that, upon specific stimuli, may act as transcriptional modifiers (Bean et al., 2008; Carrasco and Hidalgo, 2006). MicroRNAs (miRNAs) are small non-coding RNAs (20-22 nt long) that post-transcriptionally control gene expression and can modulate metabolism (Dumortier et al., 2013). miRNAs were also recently implicated in the regulation of mitochondrial metabolism in skeletal muscle. It was demonstrated that mice lacking miR-378-3p and −5p encoded by the PGC1beta locus are resistant to obesity induced by high-fat diet and can efficiently oxidize FAs (Carrer et al., 2012). However, whether such a regulation occurs normally and is involved in the specification of the metabolic preference of individual fibers is unknown.

To define networks of coding and non-coding RNAs that regulate skeletal muscle metabolism, a genomic approach in single isolated myofibers is required, to avoid the averaging effect of multiple fiber types and signals of non-muscle cells (Chemello et al., 2011; Chemello et al., 2015; Mammucari et al., 2015; Murgia et al., 2015). Here, we compiled a comprehensive catalogue of mRNAs and miRNAs expressed in single myofiber types of mouse muscles. This allowed us to obtain the RNA regulatory networks supervising the metabolic phenotypes of myofibers and to identify two specific miRNAs (miR-27a-3p and miR-142-3p) that, by influencing mitochondrial shape and respiration, define lipid and glycogen utilization and hence the oxidative and glycolytic myofiber traits.

## RESULTS

### The transcriptional signature of a myofiber echoes its metabolic trait

We undertook a large-scale expression profiling of single isolated myofibers, by collecting 100 myofibers from *soleus* and 100 myofibers from EDL mouse muscles. In a fragment of each fiber, we measured MyHC isoforms as a reference to the canonical fiber catalogue (**Figure 1A and S1**). As expected, this method identified 6 major patterns of MyHC isoforms, namely 1, 2A, 2A/2X, 2X for *soleus*, and 2X/2B, 2B for EDL (**Table S1**). We therefore profiled 10 myofibers from each of the six major MyHC groups (Dataset S1), generating a collection of 60 transcriptional profiles. Totally, 1,936 probes resulted differentially expressed (DE) among the fiber repertoire (Dataset S2). Cluster analysis gathered the 60 profiles in a larger signature composed by 40 fibers and in a smaller one summing up the remaining 20 fibers (**Figure 1B**). The larger signature was clearly divided in two sub-signatures. One, that we named transcriptional slow (tS, 14 *soleus* fibers), was mainly composed by type 1 fibers and the other, called transcriptional intermediate (tI, 26 *soleus* fibers), was composed by type 2A, 2A/2X and 2X fibers. The smaller signature, composed by type 2B and 2X/2B fibers, was named transcriptional fast (tF, 20 fibers from EDL). To obtain a quick and univocal assay for the assignment of myofibers to one of the three signatures, we identified the 5 genes with the best classification power (**Figure S2A**), further including Myh2 transcript as the best marker for tI signature (**Figure S2B and Dataset S3**). As for MyHC proteins, at transcriptional level Myh mRNAs are excellent markers for myofiber typing (**Figure S2C**).

**Figure 1:**
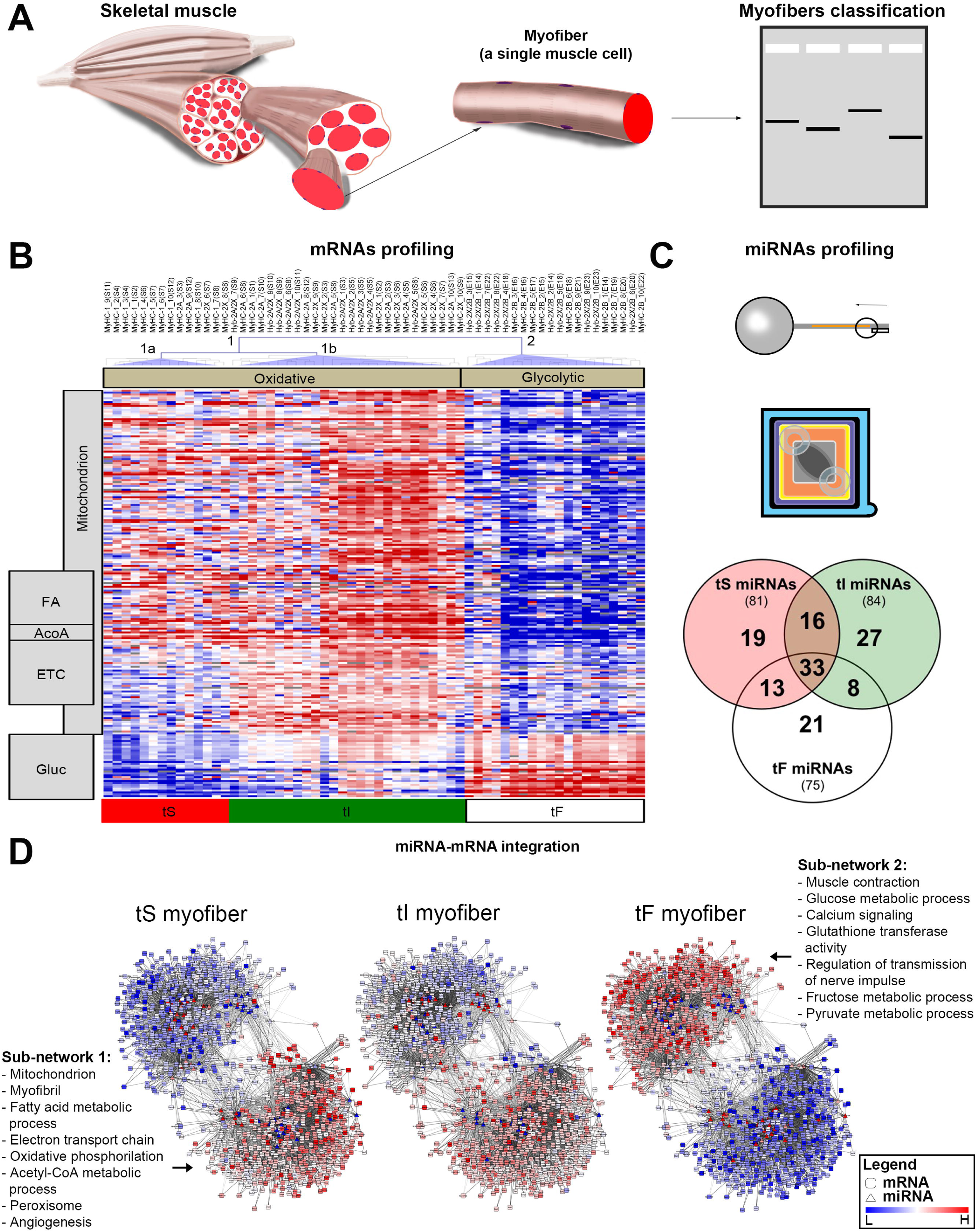
Transcriptional signatures of myofibers from *soleus* and EDL mouse muscles. (A) Single isolated myofibers were dissociated from *soleus* and EDL mouse muscles and classified by MyHC isoform pattern. (B) Cluster analysis of the DE microarray probes identifies two main signatures (1 and 2) that reflect fiber metabolism (oxidative and glycolytic). Oxidative signature (1) is further divided in two groups (1a and 1b) that we named transcriptional slow (tS) and intermediate (tI). MyHC isoform composition was indicated for pure (MyHC) or hybrid (Hyb) types. Myofibers of the same type were numbered from 1 to 10 and marked with the muscle of origin (S for *soleus*, and E for EDL) and the mouse number (parentheses). The heat-map shows the expression of DE genes involved in metabolism as identified by GO analysis (**Table 1**) (Blue is for low expression, red for high and grey for under detection limit). On the left are indicated gene functions: FA metabolic process (FA), acetyl-CoA metabolic process (AcoA), electron transport chain (ETC) and glucose metabolism (Gluc). (C) miRNA libraries were constructed and amplified using the SMART technology and profiled by NGS. A total of 137 miRNAs expressed in at least one myofiber transcriptional type was identified: 81 miRNAs were found more expressed in tS myofibers, 84 in tI myofibers, and 75 in tF myofibers. 33 miRNAs (including the myomiRs miR-1a-3p and −133a-3p) are generally expressed across all myofibers. (D) Networks of post-transcriptional regulation in tS, tI and tF myofibers. Triangles are for miRNAs and squares for mRNAs, while colors reflect their expression levels). GO enrichment analysis was performed on genes of the two sub-networks, and only terms with an enrichment score > 1.3 are listed (the highest at the top). For larger images see **Figure S5**.

To understand the functional meaning of the gene clustering architecture, which divided the DE probes in 8 groups (**Figure S2D and Dataset S4**), we categorized them evidencing that the biological processes differing between the 3 myofiber signatures are principally related to cellular metabolism (**Table 1 and Dataset S5**). In fact, the two major clusters of fiber profiles reflected the two metabolic traits of skeletal muscle tissue, i.e. oxidative and glycolytic. tS and tI myofibers were enriched in genes coding for enzymes involved in FA, electron transport chain, and acetyl-CoA metabolic processes (Figure 1B); tF myofibers expressed preferentially genes coding for glucose metabolism proteins (Figure 1B). These experiments confirmed that different myofiber types display specific metabolic gene signatures. Interestingly, Gene Ontology analysis indicated that mitochondrial genes are up-regulated especially in tI myofibers, suggesting the high importance of the organelle in the metabolism of this type of myofibers.

**Table 1:** Gene Ontology analysis of genes differentially expressed among transcriptional myofiber types. Gene Ontology (GO) enrichment analysis of the 1,936 DE microarray probes was separately performed on genes that share common expression patterns among the 3 transcriptional myofiber signatures (tS, tI, and tF) using DAVID bioinformatic tool. One representative GO term from each functional annotation cluster with an enrichment score cutoff > 1.3 (the higher, the better) and P-values < 0.05 (the lower, the better) was listed. Overlapping functional terms were ignored and only groups from the top 5 clusters are shown (additional information in **Dataset S5**).

|  | Enrichment Score | P-value |
| --- | --- | --- |
| **Probes overexpressed in tS myofibers (cluster 1, 114 probes, 94 DAVID IDs)** |  |  |
| Muscle tissue development | 2.14 | 3.03E-04 |
| Cytoskeletal protein binding | 1.96 | 4.57E-04 |
| **Probes overexpressed in tS and tI myofibers (clusters 2 and 3, 806 probes, 643 DAVID IDs)** |  |  |
| Mitochondrion | 23.80 | 9.37E-33 |
| Fatty acid metabolic process | 8.51 | 6.12E-12 |
| Myofibril | 8.18 | 6.95E-10 |
| Electron transport chain | 4.65 | 5.71E-06 |
| Acetyl-CoA metabolic process | 3.52 | 1.87E-07 |
| **Probes overexpressed in tI myofibers (cluster 4, 95 probes, 85 DAVID IDs)** |  |  |
| Mitochondrion | 7.95 | 2.40E-18 |
| Electron transport chain | 5.63 | 4.12E-13 |
| **Probes overexpressed in tI and tF myofibers (cluster 5, 240 probes, 123 DAVID IDs)** |  |  |
| Glucose metabolic process | 9.23 | 6.64E-10 |
| Muscle contraction | 2.40 | 2.57E-03 |
| Myofibril | 1.98 | 8.80E-04 |
| **Probes overexpressed in tF myofibers (clusters 6 and 7, 638 probes, 506 DAVID IDs)** |  |  |
| Sarcoplasmic reticulum | 8.06 | 8.06E-11 |
| Myofibril | 5.61 | 6.27E-07 |
| Glutathione transferase activity | 3.16 | 2.13E-03 |
| Glucose metabolic process | 2.90 | 1.59E-05 |
| Muscle contraction | 2.79 | 3.73E-04 |

### MicroRNAs post-transcriptionally regulate myofiber metabolism

miRNAs are important post-transcriptional regulators of cell metabolic pathways (Dumortier et al., 2013; Rottiers and Naar, 2012). We explored their role in the regulation of myofiber metabolic mRNA signatures. Populations of mRNAs and miRNAs were purified and separated from single isolated myofibers. The miRNA fractions were analysed by next-generation sequencing (NGS) in each fiber type, transcriptionally classified using the previously established mRNA markers (**Figure S3**). We identified 81 miRNAs expressed in tS, 84 in tI, and 75 in tF myofibers (**Figure 1C** and **Dataset S6**). We recognized 15 significantly enriched families, belonging to known muscle-specific miRNAs (myomiRs; (McCarthy, 2011), namely miR-1a, b/206/613, miR-133a, b, c and miR-486-5p/3107; **Dataset S7**). In association to miR-1a-3p and miR-133a-3p (which we confirmed as two of the most expressed miRNAs in muscle fibers) we identified 10 other miRNAs highly expressed in all fiber types (**Table S2**). These include almost all members of the let-7 family, indicating their importance in skeletal muscle biology. Interestingly, we could associate specific miRNAs to different transcriptional signatures (**Figure S4A**). For example, myomiR-206-3p and miR-208b-3p that control skeletal muscle oxidative metabolism (Boettger et al., 2014; Gan et al., 2013) are strictly expressed in tS myofibers. Quantitative RT-PCR measures of 55 miRNAs confirmed the NGS results with a correlation of 0.78 (**Figure S4B** and **Dataset S8**). Certain miRNA-3p/miRNA-5p couples were differentially expressed across different myofibers, as already evidenced in other tissue types (Martini et al., 2014).

The identification of miRNA-targets in an mRNA population is critical. For this reason, putative miRNA-targets were extracted from the repertoire of mRNAs expressed in muscle fibers by predictive algorithms and filtered for biological significance. As miRNAs act through target degradation (Hu and Coller, 2012), only miRNA-mRNA groups that showed a negative correlation between their expression levels were selected for the construction of an integrated network. A total of 5,968 interactions were computed, resulting from the negative correlations between the expression of 79 miRNAs and 1,051 putative mRNA targets (**Dataset S9**). The topology of the miRNA-mRNA interactions clearly draws two sub-networks where miRNAs are placed as central regulators (**Figure 1D** and **Figure S5**). The sub-network 1 is mainly characterized by genes coding for mitochondrial proteins and enzymes involved in oxidative phosphorylation and FA metabolism, whereas the sub-network 2 contained genes encoding for glycolytic enzymes. Considering the number of edges for each node in the network it follows that each miRNA, on average, contacts 81 mRNAs, supporting the idea that a single miRNA may regulate many mRNAs. Comparing the networks for tS and tF myofibers an opposite expression trend is clearly observed for connected genes and miRNAs (**Figure 2D**). In tS fibers, transcripts involved in glycolytic metabolism were generally under-expressed while the connected miRNAs were up-regulated; conversely, transcripts involved in oxidative phosphorylation and FA metabolism were up-regulated, while the connected miRNAs were down-regulated. The picture was specular in tF fibers. The network for tI fibers generally showed intermediate expression levels. In summary, this set of data provides the first cell-specific expression network that may control the glycolytic and oxidative metabolic traits.

**Figure 2:**
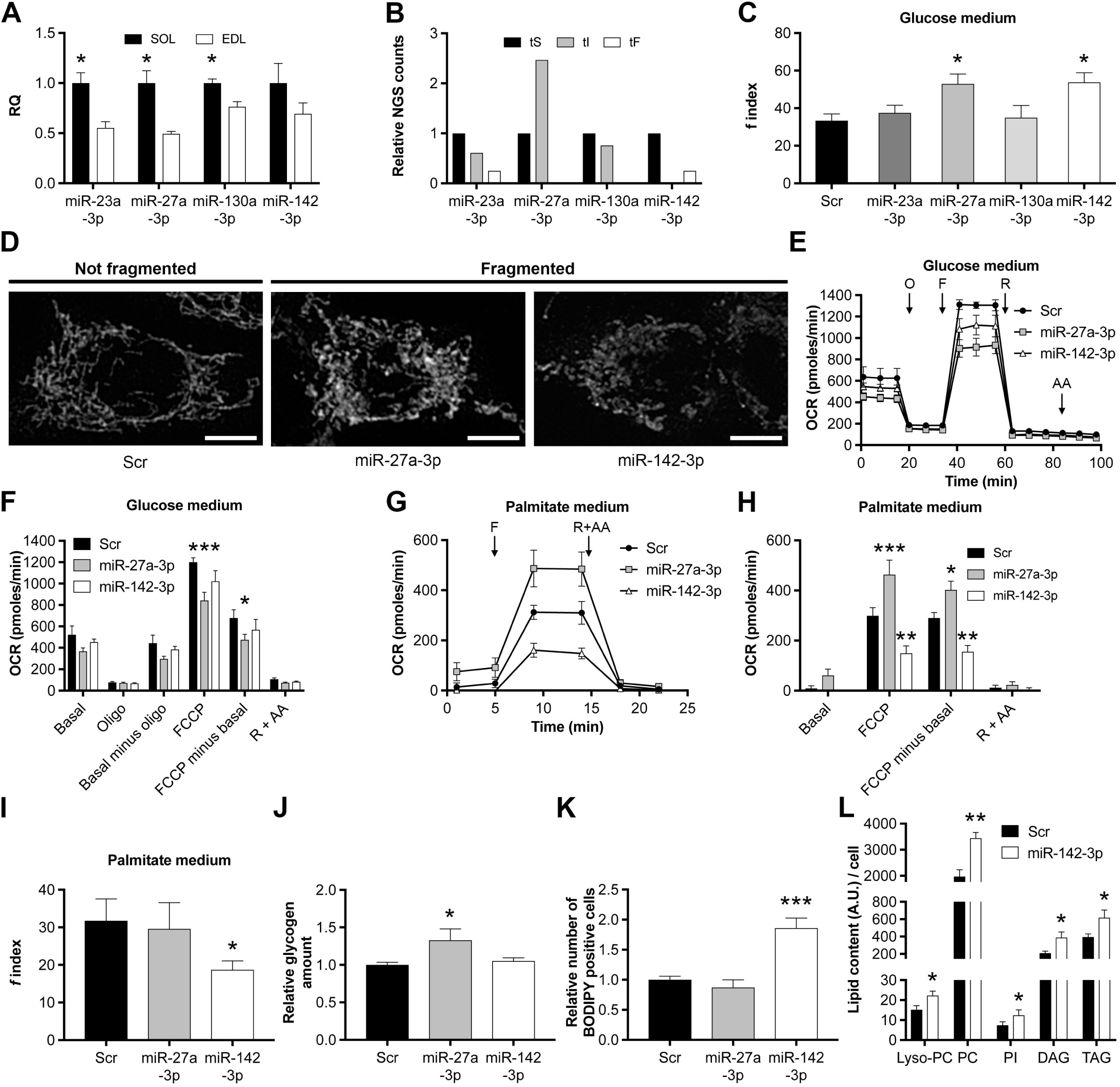
Effects of miR-27a-3p and −142-3p on mitochondrial morphology and energy metabolism: *in vitro* studies. (A) Relative quantification (RQ) by qRT-PCR of the selected miRNAs. miRNA expression levels were tested in a slow muscle (*soleus*, SOL) and in a fast muscle (EDL, used for data normalization). Each experiment was performed at least in triplicate. (B) NGS results for the selected miRNAs on tS, tI and tF myofibers. Expression levels were normalized on tS myofibers. (C) Quantification of the fragmentation index *f* of mitochondrial networks after miRNA transfections. *n* = at least 20 for each experiment. (D) Representative fluorescence microscopy images of single cells with different levels of mitochondria fragmentation. Mitochondria were stained with anti-TOM20 and imaged by confocal microscopy. Images of each cell were captured at different focal depths and then processed by focal plane merging and convolution. Scale bar is 10 µm. (E) and (G) Bioenergetic profiles of transfected myoblasts in glucose or palmitate substrate medium. OCR of C2C12 myoblasts transfected with miRNA mimics was evaluated with the sensitive Seahorse technology after 72 hours of transfection. Bioenergetic profiles of cells were measured in a basal state and after the addition of oligomycin (O, to inhibit ATP synthase), FCCP (F, to uncouple ATP synthesis from the electron transport chain), and rotenone and antimycin A (R and AA, to block respectively complexes I and III of the electron transport chain). In palmitate medium, lipid specific respiration was analysed by acute Cpt1 inhibition by etomoxir. miRNA transfections were independently replicates at least three times, and for each one five technical replicates were measured. (F) and (H) Parameters of mitochondrial function of transfected myoblasts in glucose or palmitate resulting from the bioenergetic profiles. (I) Quantification of the fragmentation index *f* of mitochondrial networks after miR-27a-3p or miR-142-3p transfections in palmitate medium. Control experiments performed with Bovine Serum Albumine alone (BSA, necessary for conjugate palmitate), confirmed that miR-142-3p mitochondrial fragmentation is induced by palmitate (data not shown). For each experiment, *n* = at least 20. (J) C2C12 myoblasts were transfected with a scramble sequence as control, with miR-27a-3p, or with miR-142-3p. Cells were lysed and glycogen content was quantified using a luminometer. Data are the mean of two replicates for three independent transfection experiments (*n* = 6). (K) Transfected myoblasts were stained for lipids with BODIPY fluorophore and fluorescence positive cells were counted by flow-cytometry. Data are the mean of two replicates for three independent transfection experiments (*n* = 6). (L) Lipids were extracted from myoblasts transfected with a control scramble sequence or with miR-142-3p, and measured with liquid chromatography-mass spectrometry (LC-MS) technology. Classes of lipids significantly different between samples are represented in the figure (Lyso-PC: lysophosphatidylcholine, PC: phosphatidylcholine, PI: phosphatidylinositol, DAG: diacylglycerol, TAG: triacylglycerol). Lipid quantification was measured as arbitrary units (A.U.) of signal areas of chromatograms and normalized to the number of cells relative to each sample. Number of independent experiments (*n*) per condition = 4. In the entire figure: Scr is the control transfected with a scrambled sequence; error bars are SEM; adjusted P-values of ANOVA statistical test were indicated as follow: * < 0.05, ** < 0.01, *** < 0.001, and they are referred in comparison to the control.

### miR-27a-3p and −142-3p influence mitochondrial shape and respiration *in vitro*

Our data indicate that differences in mitochondrial genes characterize the myofiber signatures, particularly those of tI myofibers, (**Figure 1B** and **Table 1**), supporting the role of this organelle in directing metabolic plasticity of skeletal muscle (Hood et al., 2006). We selected for functional analyses a set of miRNAs that are connected though a different number of transcripts in the metabolic networks represented in **Figure 1D**. These are miR-27a-3p and −23a-3p, connected to 114 and 105 putative mRNA targets; and miR-130a-3p and −142-3p connected to 73 and 48 mRNA targets. In whole muscle analysis, all these miRNAs are more expressed in slow muscle, except for miR-142-3p that does not vary between fast and slow muscles (**Figure 2A**). NGS analysis on single myofibers revealed that miR-23a-3p and −130a-3p were up-regulated in tS and tI myofibers, miR-27a-3p was up-regulated in tI myofibers and, on the contrary, miR-142-3p was down-regulated in tI myofibers (**Figure 2B**). Since function and shape of mitochondria are strictly related (Schrepfer and Scorrano, 2016), we examined the role of the four selected miRNAs in mitochondrial morphology. In mouse muscle C2C12 cells, the modification of miR-23a-3p or miR-130a-3p expression did not affect mitochondrial shape, while increased levels of miR-27a-3p or miR-142-3p caused mitochondrial fragmentation (**Figure 2, C and D**). Electron microscopy confirmed that myoblasts transfected with these two miRNAs possess smaller and less-connected mitochondria, although with normal cristae structure (**Figure S6A**). The altered mitochondrial morphology was accompanied by changes in mitochondrial fuel oxidation. In myoblasts relying on glucose as oxidable substrate, miR-27a-3p caused a significant decrease in maximal oxygen consumption rate (OCR) induced by FCCP (**Figure 2, E and F**). When the oxidable substrate was palmitate, miR-142-3p decreased, whereas miR-27a-3p increased cellular respiration (**Figure 2, G and H**). Coherently, myoblasts over-expressing miR-142-3p showed mitochondrial fragmentation in palmitate medium and altered cristae structure, whereas no alteration in cristae structure was observed in myoblasts over-expressing miR-27a-3p (**Figures 2I, S6A, and S6B**). We investigated if the alteration of bioenergetics supply, observed in myoblasts transfected *in vitro* with miR-27a-3p or −142-3p, could result in accumulation of unused nutrients that eventually leads to the observed mitochondrial fragmentation, as already demonstrated in other experimental paradigms (Liesa and Shirihai, 2013; Schrepfer and Scorrano, 2016). miR-27a-3p induced the accumulation of glycogen (**Figure 2J**), whereas miR-142-3p that of lipid droplets in myoblasts (**Figure 2K**), in the proximity of mitochondria (**Figure S6C**). To understand what type of lipids were accumulated in myoblasts transfected with miR-142-3p, we profiled them by lipidomics. This unbiased analysis distinguished 66 lipids, belonging to 10 different classes (**Dataset S10**) and confirmed an overall increase in lipid content after miR-142-3p over-expression (**Figure S6D** and **S6E** for the classes not significantly altered). Phosphatidylcholine (the major phospholipid populating the surface of lipid droplets) and triacylglycerol (the main constituent of the neutral lipid core of lipid droplets) are among the classes of lipids that increased (**Figure 2L**), being probably the major components of miR-142-3p induced fat droplets.

These results indicate that miR-27a-3p and miR-142-3p act on cellular metabolic pathways inducing morphological and physiological dysfunction of mitochondria. FA utilization is inhibited by miR-142-3p, and it is improved by miR-27a-3p, whereas glycogen consumption is limited by over-expression of miR-27a-3p. Gene expression analyses of myoblasts over-expressing miR-27a-3p or miR-142-3p were coherent with these functional changes. In fact, miR-27a-3p depressed electron transport chain genes, whereas miR-142-3p turned off genes coding for enzymes involved in FA metabolism (**Dataset S11**).

### miR-27a-3p and −142-3p influence metabolism of muscle cells by targeting Pgm2, Gaa and Fndc5

We next evaluated if miR-27a-3p and miR-142-3p down-regulated the putative targets identified by the integration of mRNA and miRNA myofibers expression signatures (**Figure 2C**). Microarray analysis showed that 66.7% and 73.9% of miR-27a-3p or miR-142-3p targets were indeed down-regulated (**Figure 3A**). Interestingly, peroxisome proliferator-activated receptor γ (Pparg), a validated target of miR-27a-3p (Kim et al., 2010) was down-regulated by 1.5 fold. Because miR-27a-3p influences glycogen utilization, we searched for genes involved in glycogenolysis in the list of putative targets down-regulated in C2C12 cells over-expressing miR-27a-3p. Phosphoglucomutase (Pgm2) and acid α-glucosidase (Gaa), genes encoding for key enzymes respectively in cytosolic and lysosomal glycogenolysis, were among the targets. The interaction between miR-27a-3p and the 3’-UTR of Pgm2 and Gaa was confirmed by luciferase assay (**Figure 3B**). In addition, glycogen phosphorylase (Pygm), another important gene for glycogenolysis not directly targeted by miR-27a-3p, was under-expressed when miR-27a-3p is up-regulated in C2C12 cells (**Table S3**).

**Figure 3:**
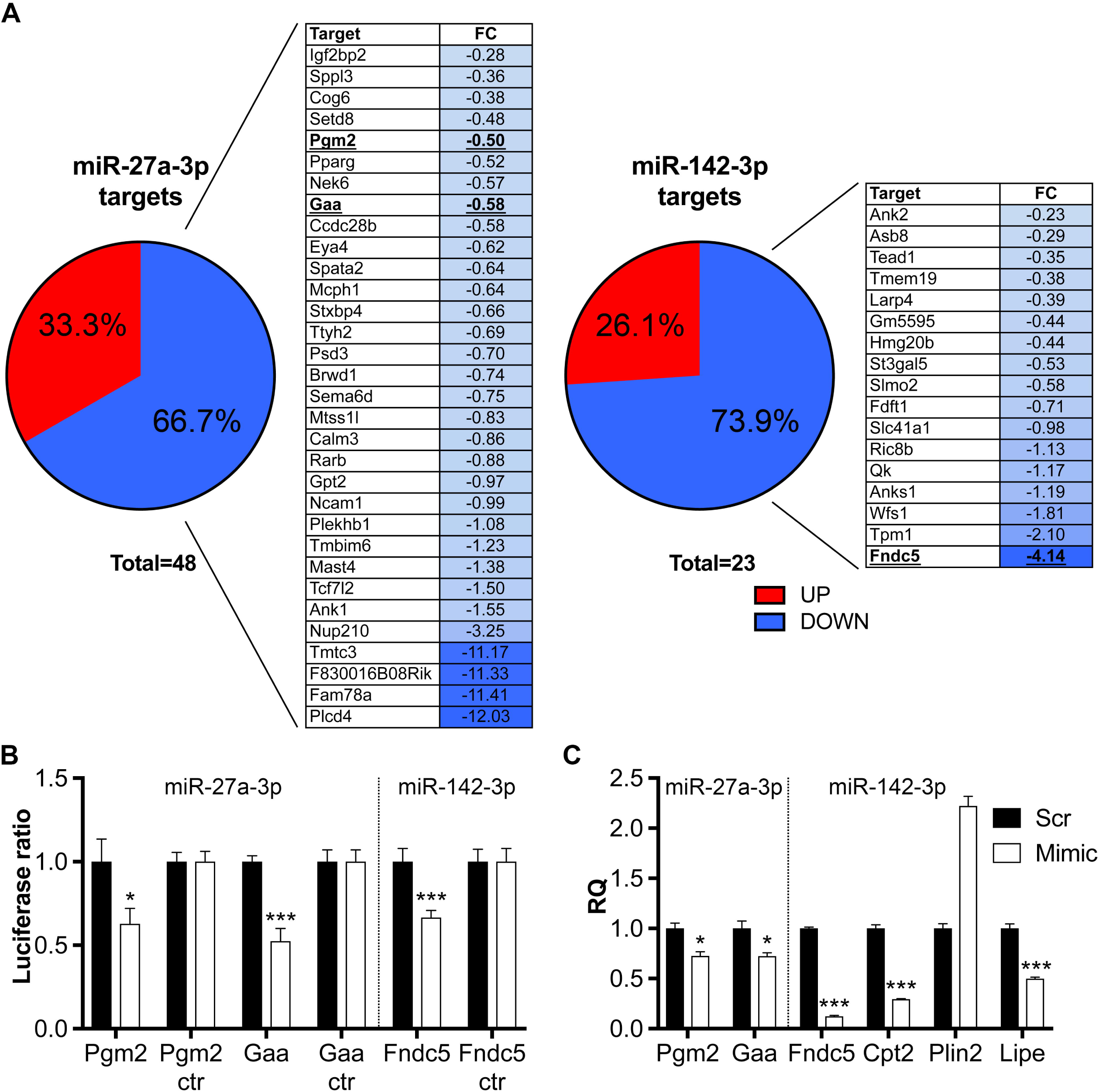
Identification and characterization of transcripts that are altered by miR-27a-3p and miR-142-3p over-expression. (A) Expression profiling of myoblasts transfected with miR-27a-3p or miR-142-3p showed that most targets identified in myofiber mRNAs-miRNAs interaction networks of Figure 2C are down-regulated. 32 transcripts out of 48 targets of miR-27a-3p (66.7%) are down-regulated in C2C12 myoblasts upon the over-expression of that miRNA. Similarly, the over-expression of miR-142-3p resulted in the down-regulation of 17 out of 23 targets (73.9%). Tables flanking pie graphs list the expression values (Log_2_ fold change “FC”) of down-regulated miRNA targets in transfected myoblasts in comparison with myoblasts transfected with a scrambled sequence as control (Scr). (B) Luciferase assays were performed to demonstrate the direct interaction between the two miRNAs and their targets. Part of the 3’-UTR sequence containing the miRNA putative interaction site (or not containing, ctr) was cloned in pmirGLO vector. Firefly luciferase (reporter gene) and renilla luciferase (control reporter for normalization) activities were measured after the transfection in C2C12 cells together with miRNA mimics or miRNA Scr sequence. Data are expressed as mean of at least five replicates; error bars are SEM; P-values of t-test are indicated as follow: * < 0.05, *** < 0.001. (C) Relative quantification (RQ) by qRT-PCR of specific transcripts, codifying for relevant metabolic enzymes, in mouse EDL muscles transfected with miR-27a-3p or −142-3p mimics. These *in vivo* experiments confirm that a) miR-27a-3p up-regulation inhibits the expression of Pgm2 and Gaa genes, and b) miR-142-3p up-regulation diminishes the expression of Fndc5 and Cpt2. Moreover, when miR-142-3p is up-regulated in muscles, Plin2, a marker of lipid droplets, is over-expressed and the lipase hormone sensitive (Lipe) is under-expressed. Data are expressed as mean of three independent transfections; error bars are SEM; P-values of t-test are indicated as follow: * < 0.05, *** < 0.001.

In C2C12 myoblasts expressing miR-142-3p, fibronectin type III domain containing 5 (Fndc5), encoding for the precursor of the myokine Irisin (Bostrom et al., 2012) was the most down-regulated gene. To validate this negative connection, we transfected C2C12 myoblasts with a construct containing the 3’-UTR of Fndc5 driving the expression of luciferase, observing a 40% reduction of activity upon over-expression of miR-142-3p (**Figure 3B**). Transcriptomic analysis of myoblasts over-expressing miR-142-3p indicated that several genes involved in FA metabolism were deregulated. Acyl-CoA synthase 1 (Acsl1), the most important isoform for the synthesis of acyl-CoA used by skeletal muscle mitochondria (Li et al., 2015), was up-regulated. This confirms an increase in FA anabolism, although other acyl-CoA synthase isoforms (Acsl3, Acsl6) were found to be down-regulated. Genes of FA catabolic enzymes, like acyl-CoA dehydrogenase (Acad10, Acad12), or related to FA catabolism, like carnitine palmitoyl transferase (Cpt2), hormone sensitive lipase (HSL), and Ucp2 and 3 were down-regulated. Conversely, the lipid droplet markers perilipins (Plin1, Plin2, Plin4) were up-regulated (**Table S3**). These *in vitro* over-expression experiments suggest the principal mediators by which miR-27a-3p and miR-142-3p might exert their differential control of fuel consumption in muscle cells.

### miR-27a-3p and −142-3p control fuel availability *in vivo*

To test whether these two miRNAs can also induce glycogen or lipid accumulation *in vivo*, we transfected hindlimb mouse muscles with miRNA mimics. In muscle over-expressing miR-27a-3p, periodic acid–Schiff (PAS) staining confirmed glycogen accumulation (**Figure 4, A and B**). Also, electron microscopy analysis established the presence of abundant glycogen granules and showed that mitochondria are impaired, but generally maintaining regular cristae structure (**Figure 4C**). The expression of miR-142-3p *in vivo* recapitulated the results obtained in C2C12 myoblasts, with stronger BODIPY staining when muscles were transfected with the miRNA mimic (**Figure 4, D and E**), and clearly visible lipid droplets in electron microscopy images (**Figure 4D**).

**Figure 4:**
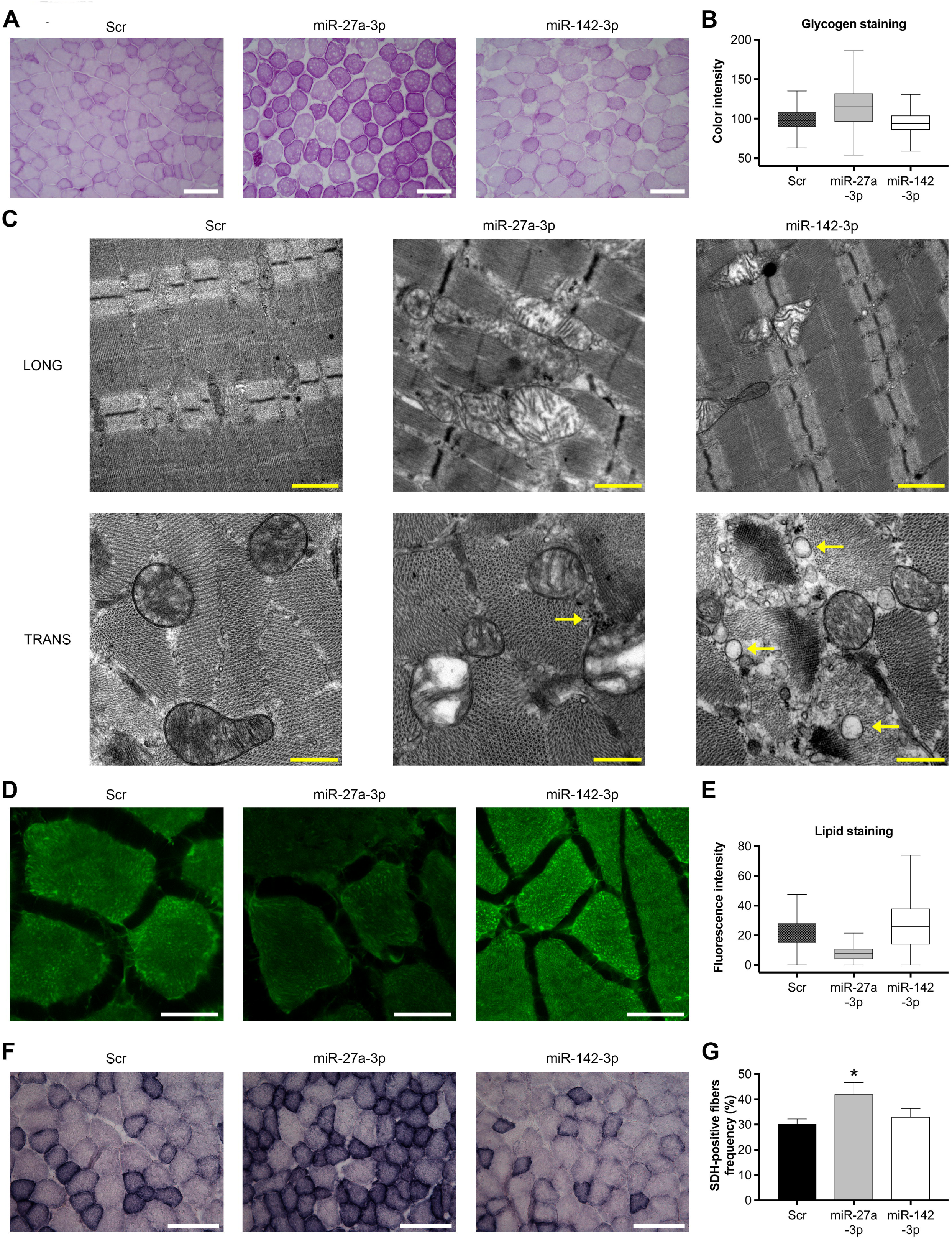
Alterations of fuel availability induced by miR-27a-3p and miR-142-3p: *in vivo* studies. (A) Representative cross sections of mouse *tibialis anterior* stained with PAS to detect glycogen, after one week of transfection with a scramble sequence as control (Scr), with miR-27a-3p, or with miR-142-3p. Scale bar is 200 µm. (B) The intensities of PAS color pixel were graphed in Tukey box-plots. Number of independent experiments (*n*) per condition = 3. (C) Electron microscopy of longitudinal (LONG) and transverse (TRANS) sections of transfected *gastrocnemius* muscle. Arrows indicate glycogen granules in muscles over-expressing miR-27a-3p, or lipid droplets in muscles over-expressing miR-142-3p. Scale bar is 500 nm. (D) Representative cross sections of mouse *tibialis anterior* stained for lipids with BODIPY, after one week of transfection with a scramble sequence as control, with miR-27a-3p, or with miR-142-3p. Scale bar is 50 µm. (E) BODIPY fluorescence pixel intensities were graphed in Tukey box-plots. Number of independent experiments (*n*) per condition = 3. (F) Representative cross section of mouse muscle *gastrocnemius* after one week of transfection. Oxidative myofibers are positive to succinate dehydrogenase (SDH) staining (blue color). Scale bar is 100 µm. (G) Quantification of SDH-positive fibers. At least 2,500 fibers were analyzed for each transfection. Error bars are SEM; adjusted P-values of ANOVA statistical test is indicated as follows: * < 0.05, and referred in comparison to the control.

Interestingly, upon miR-27a-3p over-expression succinate dehydrogenase staining indicated an increase in oxidative myofibers and a reduction in glycolytic myofibers, in agreement with the over-expression of mRNAs for oxidative enzymes in tI myofibers. No such variations were observed instead in muscles transfected with miR-142-3p (**Figure 4, F and G**).

qRT-PCR experiments on transfected muscles confirmed the reduced expression of the same miRNA targets already established with the *in vitro* experiments (**Figure 3C**). Pgm2 and Gaa expression was inhibited by miR-27a-3p, whereas Fndc5 was down-regulated by miR-142-3p. Furthermore, other master genes involved in FA metabolism such as *Cpt2*, *Plin2*, and *Lipe* were down-regulated in muscles upon miR-142-3p transfection.

In summary, all the results of *in vitro* and *in vivo* experiments agreed in showing the differential role of miR-27a-3p and −142-3p in the regulation of fuel consumption in skeletal muscle.

## DISCUSSION

In this work, we describe how the metabolic plasticity of myofibers relies on dynamic transcriptional networks that govern their fuel preferences and, by generating and browsing the compendium of mRNAs and miRNAs expressed in single isolated myofibers, we identify two miRNAs that can modulate the lipid utilization in myofibers.

Previous transcriptional catalogues obtained from whole muscles samples (Campbell et al., 2001; Raz et al., 2018; Wu et al., 2003) represent an average transcriptome of the different muscle cell repertoires, and lack the resolution power to identify the specific signatures of the different myofiber types. This is particularly important when inspecting the transcriptional signatures that dictate the metabolic traits. However, the evolution of single cell approaches allows the identification of signatures that specify functional and metabolic behaviour of the individual myofiber type. Our analysis of transcriptional signatures shows that the preferential metabolism (oxidative or glycolytic) is the key component that describes the myofiber molecular phenotype. tS and tI myofibers display high expression levels of genes encoding for enzymes of oxidative phosphorylation and FA oxidation, whereas tF myofibers preferentially express genes encoding for enzymes of glycolytic metabolism. This picture is closer to the more traditional functional classification of muscle myofibers, that differentiates them in slow-oxidative, fast-oxidative, and fast-glycolytic (Peter JB et al., 1972), than to the MyHC-based classification. Interestingly, we observed that tI myofibers (mostly composed by type 2A, 2X, and 2A/2X myofibers) are characterized by the highest level of expression of genes coding for mitochondrial enzymes. This observation agrees with previous proteomic analysis of single myofibers (Murgia M et al., 2015) that found an enrichment of mitochondrial proteins involved in oxidative phosphorylation in type 2A fibers, in accordance with their largest content in mitochondria (Schiaffino S et al., 1970).

Microarrays and NGS are approaches particularly suitable to snapshot the transcriptome of single myofibers; yet, to profile non-coding RNAs and to assign them to a specific myofiber subtype, miRNA and mRNA expression profiles must be integrated. Such an approach identified post-transcriptional circuits involved in the regulation of myofiber metabolism, in addition to those already established involving miR-208b-3p, miR-499-5p and Sox6 (Gan et al., 2013; van Rooij et al., 2009). Our integrated approach highlights two previously unappreciated major circuits that connect metabolic transcripts with miRNAs 27a-3p and 142-3p. miRNA profiling in whole muscle described these miRNAs as more expressed in slow muscle or not differentially expressed between fast and slow muscle, respectively. The single myofiber approach instead revealed their opposite expression in the tI myofibers, miR-27a-3p being highly, and miR-142-3p poorly expressed. This set of findings suggests a complementary effect of these two miRNAs in regulating myofiber metabolism. Functional assays *in vitro* and *in vivo* reveal that miR-27a-3p improves lipid utilization and reduces the breakdown of glycogen and consequently the glycolysis (**Figure 5A**). These adaptations in the selection of specific fuel type due to miR-27a-3p are coherent with the oxidative metabolism of tI myofibers, and explain the ability of the miRNA to change the myofiber phenotype from glycolytic to oxidative. On the contrary, miR-142-3p inhibits lipid utilization in myofibers (**Figure 5A**). However, also tS myofibers, which use lipids as fuel, express miR-142-3p suggesting the different repertoire of miRNA expression in tS and tI fibers can differently influence genes involved in lipid catabolism.

**Figure 5:**
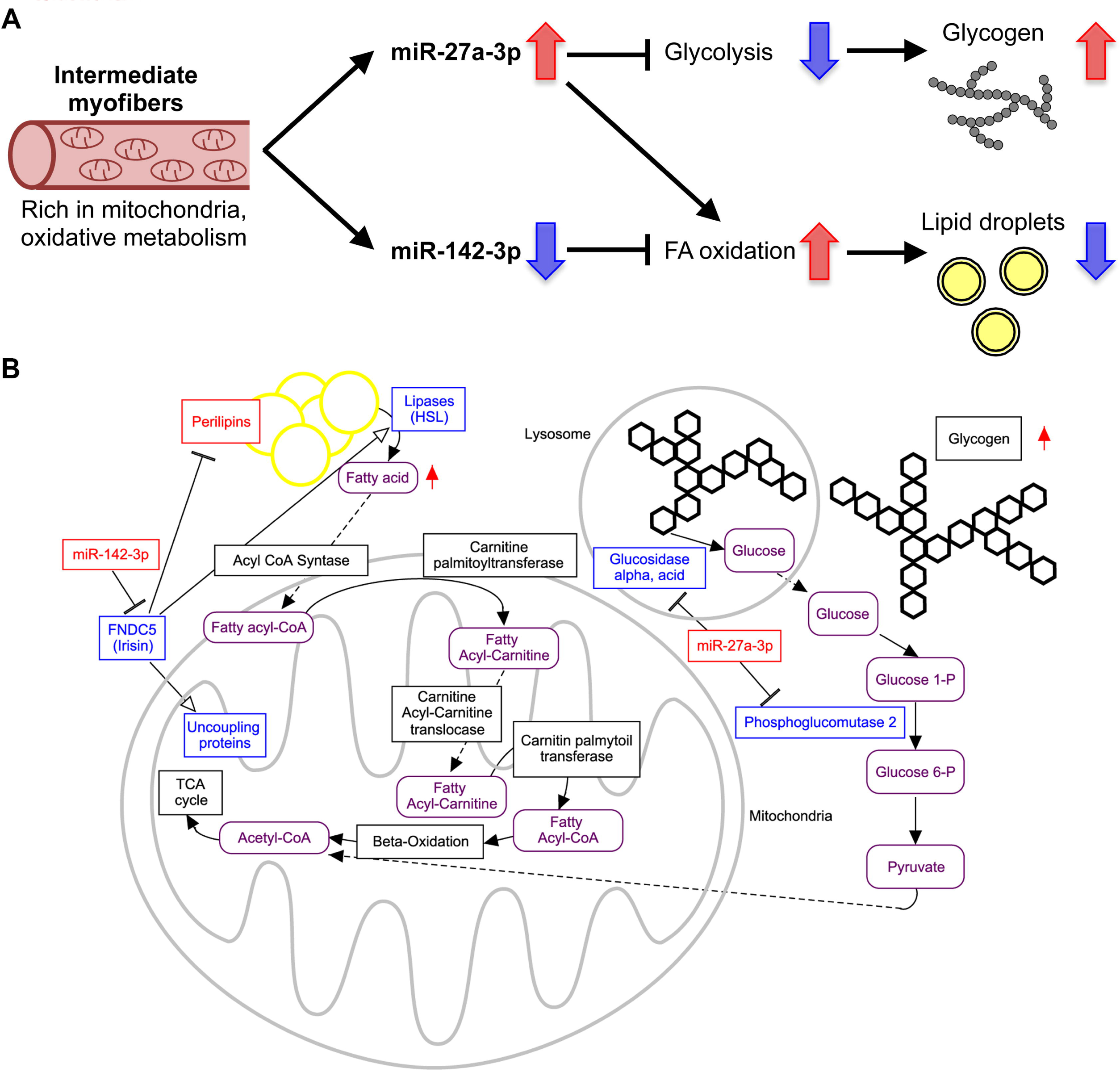
Role of miR-27a-3p and −142-3p in myofiber metabolism. (A) A model that integrates the circuits by which miR-27a-3p and −142-3p may orchestrate glycogen and FA metabolism in tI myofibers. (B) A more-detailed scheme representing miRNA modulation of myofiber metabolism. For all the components of the scheme, blue color indicates low levels and red color high levels. Asterisks indicate the miRNAs of interest.

Increased miRNAs dosage leads to accumulation of unused nutrients (i.e., glycogen and lipids) and to mitochondrial fragmentation, which appears to be due to fission. This event is likely to be secondary to the accumulation of reduced nutrients, as opposed to the elongation occurring upon nutrient starvation (Gomes et al., 2011; Rambold et al., 2011). While changes in morphology appear ancillary, mitochondrial fuel utilization is conversely embedded in the program controlled by these metabolic signatures. Expression of miR-27a-3p results in reduced maximal mitochondrial respiration, and in glycogen accumulation due to inhibition of cytosolic and lysosomal glycogen breakdown. Indeed, we found that miR-27a-3p expression represses Pgm2 (an enzyme involved in the conversion of glucose 1 phosphate generated by the glycogen phosphorylase into glucose 6 phosphate) and Gaa (a lysosomal enzyme essential for the hydrolysis of glycogen to glucose) (**Figure 5B**). Concomitantly, miR-27a-3p improves lipids utilization in agreement with its established role in adipocytes and hepatocytes (Ji et al., 2009; Shirasaki et al., 2013; Wang et al., 2011).

miR-142-3p impairs FA utilization, resulting in the accumulation of lipid droplets. Indeed, miR-142-3p expression is lowest in tI myofibers, characterized by the highest expression of mitochondrial genes. Conversely, miR-142-3p over-expression shuts down the FA mitochondrial transporter carnitine palmitoyltransferase and the uncoupling proteins Ucp2 and Ucp3. Ucp3 prevents triglyceride storage (Musa et al., 2012) and its expression increases in C2C12 myoblasts exposed to the myokine irisin (Vaughan et al., 2014). Interestingly, the irisin precursor Fndc5 is the most down-regulated among miR-142-3p targets (**Figure 5B**). miR-142-3p therefore emerges as a crucial molecule controlling the Fndc5/irisin pathway since it inhibits irisin autocrine effects, impairing FA utilization and eventually leading to lipid droplets accumulation. Indeed, miR-142-3p down-regulates HSL and up-regulates perilipins, whereas Fndc5/irisin promotes lipolysis exactly by triggering the opposite effect on HSL and perilipins (Xiong et al., 2015).

The miRNAs and targets discovered in our work are likely participants in obesity and diabetes. Indeed, a rat model of type 2 diabetes and mature adipocytes from obese mice display deregulation of miR-27a-3p as well as of its target Gpd2 (Herrera et al., 2010; Kim et al., 2010; MacDonald et al., 1996). miR-142-3p was associated with alterations in lipid pathways and abnormal fat deposition also in brain (Di Paolo and Kim, 2011) and it is a biomarker for morbidity of obesity and type 2 diabetes (Ortega et al., 2013; Ortega et al., 2014). We can envisage that a muscular metabolic reprogramming by delivery of miRNAs or antagomirs detailed in this work could be a promising route for the development of treatments counteracting the pathological side effects of diabetes and obesity.

## ACKNOWLEDGMENTS

Authors wish to thank dr. Federico Caicci (BioImaging Facility of the Department of Biology, University of Padova) for help with electron microscopy analyses; dr. Caterina Millino and dr. Beniamina Pacchioni (Microarray Service MicroCribi, CRIBI Biotechnology Centre, University of Padova), for the help with microarray analyses; Professor Fabio Di Lisa and dr. Valeria Petronilli for useful discussions. This work was supported by grants of the CARIPARO Foundation (FIBRE-GEXP) and of the University of Padova (CPDA139317) to S.C., Cariplo foundation (MAYBE) to S.C. and G.L., by FP7 CIG PCIG13-GA-2013-618697, Italian Ministry of Research FIRB RBAP11Z3YA_005, and by EFSD/Novo Nordisk Programme for Diabetes Research in Europe grant to L.S. F.G. and E.H-C. were supported by a DTI-IMPORT FP7 Fellowship.

## AUTHOR CONTRIBUTIONS

G.L., C.R., L.S., and S.C. conceived the project. F.C., F.G., A.Z., P.C., E.H.C., P.M, C.B, E.A., and R.F. performed experiments. F.C., P.L., G.G., C.R., P.B., L.S., S.C. and G.L. analysed results. F.C., S.C., L.S., P.B. and G.L. wrote the manuscript, which all Authors reviewed.

## TABLES

**Table S1:** Results of myofiber typing. Single fibers were enzymatically dissociated from *soleus* (n = 13) and EDL (n = 9) mouse muscles. A small fragment of each isolated myofiber was characterized on the basis of MyHC isoform protein expression determined by SDS-PAGE. The distribution of fiber types mirrors the properties of the two muscles analyzed. All major adult fiber types were identified together with the most frequent hybrid combinations. * These myofibers, due to not typical MyHC composition, were not processed for expression profiling.

|  | **1** | **2A** | **2A/2X** | **2X** | **2X/2B** | **2B** | **Others\*** | **Total** |
| --- | --- | --- | --- | --- | --- | --- | --- | --- |
| **Soleus** | 46 | 11 | 15 | 21 | 1 | 1 | 5<br>(1/2A: 2, 1/2X: 1,<br>2X/2B: 1, 2B: 1) | 100 |
| **EDL** | 0 | 1 | 0 | 2 | 29 | 64 | 4<br>(1/2B: 1, 2A: 1,<br>2A/2B: 1,<br>2A/2X/2B: 1) | 100 |

**Table S2:**
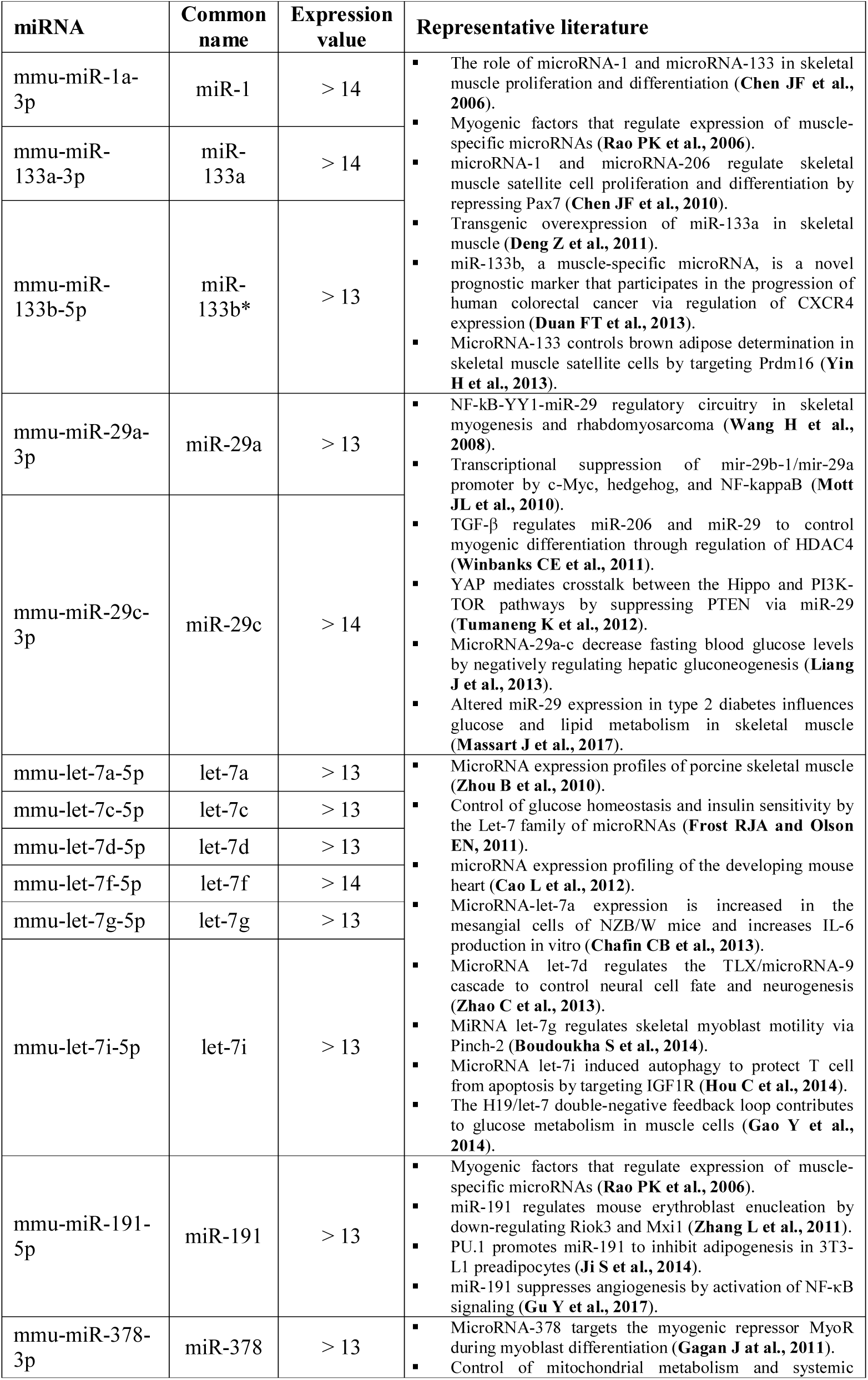
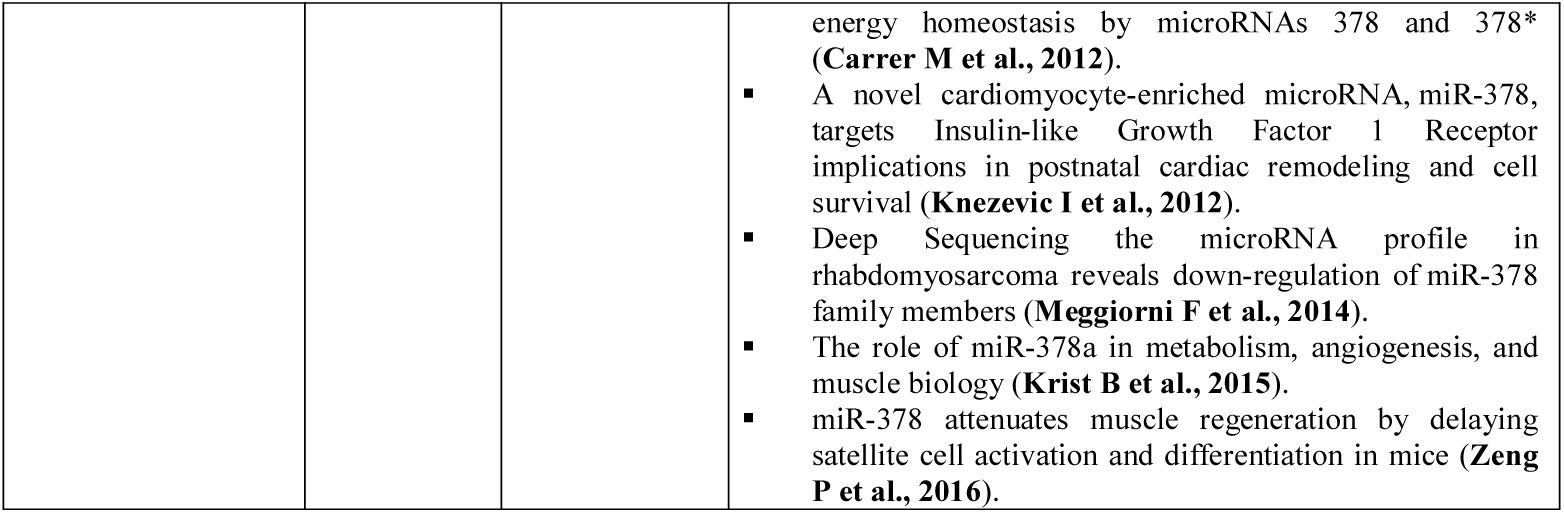
miRNAs most expressed in skeletal muscle fibers. NGS experiments performed on single isolated myofibers identified a list of 13 miRNAs highly expressed. Expression values are Log_2_ of sequencing results normalized as counts of reads per million (+1) and refer to all the three transcriptional signatures. For each miRNA or miRNA family, a list of representative literature is cited in order to elucidate their role in skeletal muscle biology.

**Table S3:**
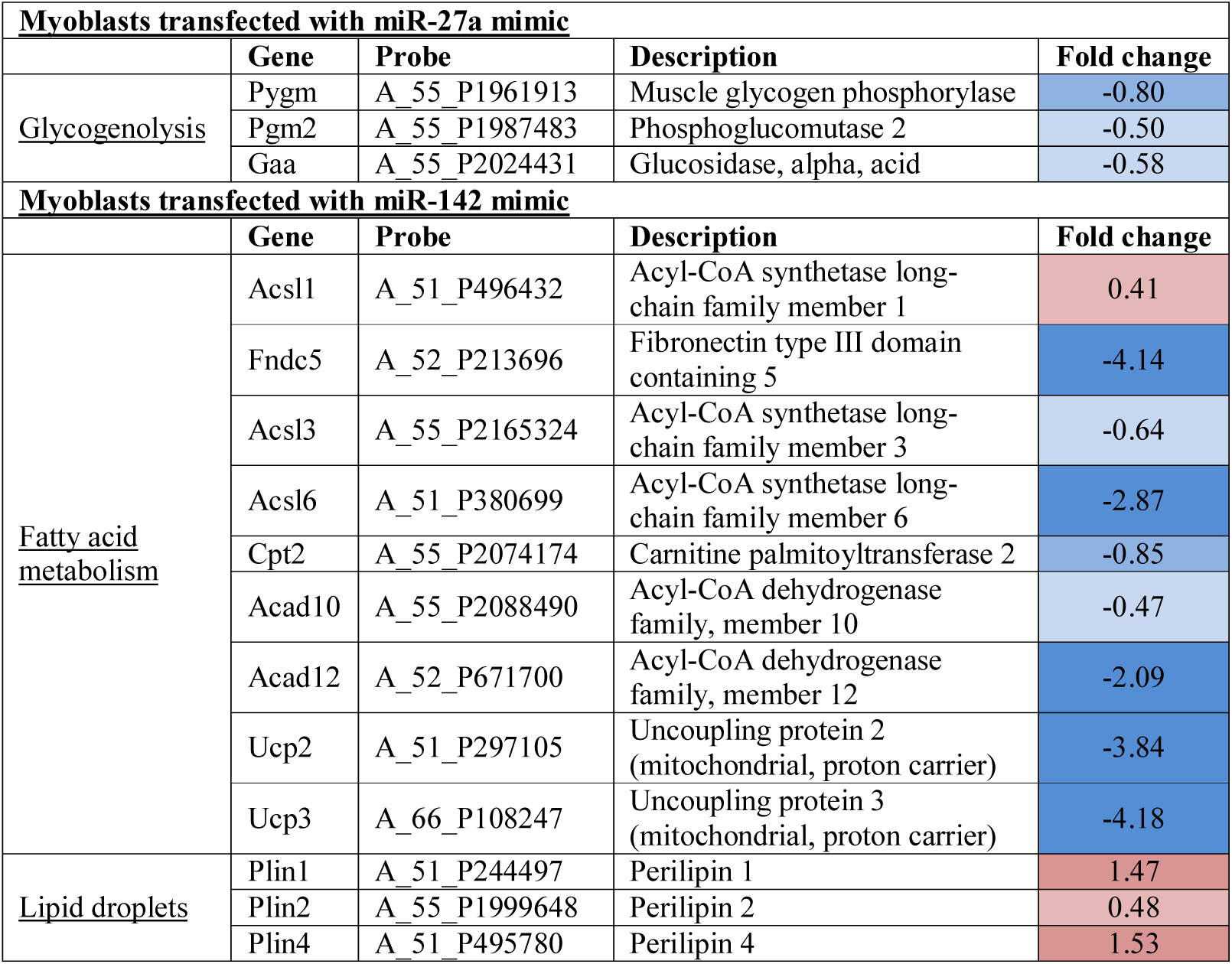
Expression changes of genes involved in the synthesis of metabolic enzymes. Expression levels of selected differentially expressed genes coding for protein involved in glycogenolysis, FA metabolism or markers of lipid droplets in myoblasts transfected with miR-27a-3p or miR-142-3p mimics. Log_2_ fold change is in comparison with myoblasts transfected with a scrambled sequence. Blue colour indicates under-expression, red colour over-expression.

**Table S4:**
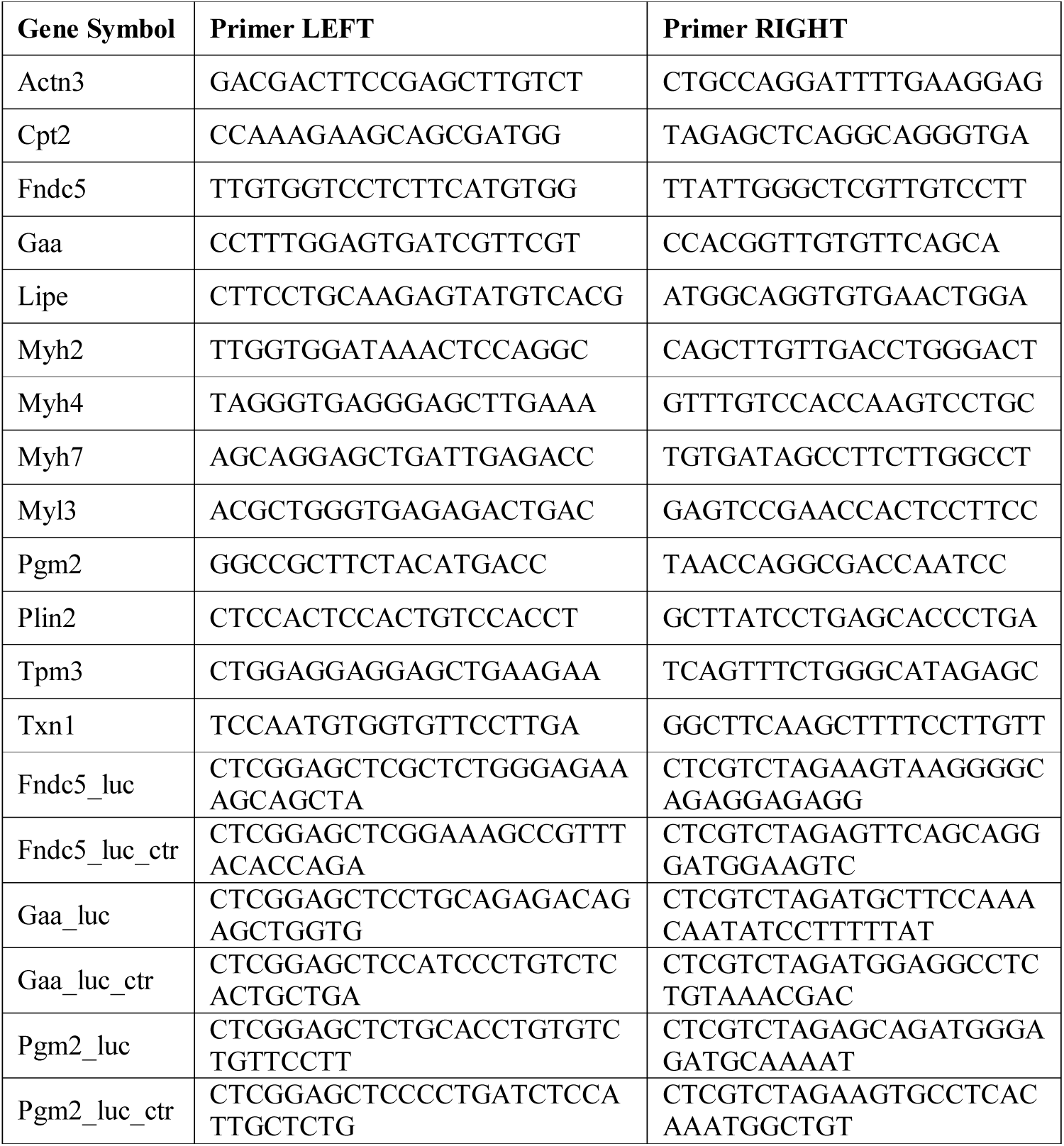
Primers used for qRT-PCRs and luciferase assays. Primers were designed using Primer3Plus.

## MATERIALS AND METHODS

### Ethic statement

All aspects of animal care and experimentation were performed in accordance with the Guide for the Care and Use of Laboratory Animals published by the National Institutes of Health (NIH Publication No. 85-23, Revised 1996) and the Italian regulations (DL 116/92) concerning the maintenance and use of laboratory animals. Experimental procedures were approved by the Ethical Committee of the University of Padova.

### Animals

Hindlimb muscles were collected from wild-type CD1 male mice at 3 months of age (Charles River, weight: 33 – 35 g). Mice were housed in a normal environment, provided with food and water and were killed by rapid cervical dislocation to minimize suffering.

### Screening of *soleus* and EDL myofibers

Enzymatic dissociation and classification of myofibers from *soleus* and EDL mouse muscles were fully described in our previous work (Chemello et al., 2011). Briefly, after collagenase treatment, each single isolated myofiber was cut and about one-third of the fiber was used for MyHC protein isoform classification by SDS-PAGE whereas the remaining part was immediately immersed in TRIzol Reagent (Thermo Fisher Scientific) for RNA purification. All myofibers were collected within 45 min. from muscle disaggregation.

### RNA isolation

Single isolated myofiber were separately lysed in 250 µl of TRIzol Reagent (Thermo Fisher Scientific) and RNA was extracted in the aqueous phase following the manufacturer’s instructions. 70% ethanol was added before column purification. For microarray experiments, total RNA was purified using the RNeasy Micro Kit (Qiagen), whereas, for miRNA analyses, separation of small and large RNAs was performed combining it with the miRNeasy Mini Kit (Qiagen) as suggested by manufacturer’s protocol. RNAs from whole muscles or cells were isolated using the standard phenol-chloroform extraction with TRIzol Reagent (Thermo Fisher Scientific) followed by isopropanol precipitation.

### Microarray profiles

#### Amplification and labeling of RNA from single fibers

In order to obtain a sufficient amount of cDNA for microarray experiments, RNA purified from a single myofiber was exponentially amplified using the TransPlex Whole Transcriptome Amplification 2 Kit (Sigma-Aldrich) in accordance with the manufacturer’s instructions. RNA was reverse-transcribed in a cDNA library, and library was amplified for 18 cycles, below the amplification “plateau” observed in a PCR test reaction. To remove the residual primers and nucleotides, amplification product was purified with the GenElute PCR Clean-up columns (Sigma-Aldrich). Resulting cDNA was quantified with Nanodrop ND-1000 spectrophotometer (Thermo Fisher Scientific). 2 μg of amplified-purified cDNA were directly labeled using the Genomic DNA Enzymatic Labeling Kit (Agilent Technologies) as described by the proprietary protocol. The kit uses random primers and the exo-Klenow fragment to directly label cDNA samples with Cy3-dUTP nucleotides. Labeled cDNA was purified using the Amicon 30kDa filters (Millipore) and quantified using NanoDrop ND-1000 spectrophotometer (Thermo Fisher Scientific). On average, cDNA yield was about 4 μg and the specific activity was of 30 pmol Cy3 per μg cDNA.

#### Amplification and labeling of RNA from cells

RNAs obtained from cells were amplified and labeled starting from 150 ng and using the Low Input Quick Amp Labeling Kit, one-color (Agilent Technologies) as described by the proprietary protocol. Labeled cRNA was purified with miRNeasy Mini Kit (Qiagen) and quantified using NanoDrop ND-1000 spectrophotometer (Thermo Fisher Scientific). On average, cRNA yield was about 10 μg and the specific activity was of 35 pmol Cy3 per μg cRNA.

#### Hybridization

Microarray experiments were performed with SurePrint G3 Mouse Gene Expression 8×60K microarray platforms (Agilent Technologies) containing 8 arrays per slide and consisting of 39,430 60mer oligonucleotide probes for Entrez Gene RNAs (GPL10787). 800 ng of labeled sample target were mixed with 5 μl of 10X Blocking Agent (Agilent Technologies) and water to a final volume of 25 μl. Samples were denatured at 95°C for 2 min. and added to 25 μl of 2X GEx Hybridization Buffer HI-RPM (Agilent Technologies). 40 μl mix was dispensed onto the array. Slides were loaded into the Agilent SureHyb chambers and hybridization was performed in a hybridization oven at 65°C for 17 hours with 10 rpm rotation. Finally, slides were washed using Wash Buffer Kit (Agilent Technologies) and dried at room temperature.

### Microarray data analysis

#### Data pre-processing

Microarray slides were scanned using G2505C scanner (Agilent Technologies) at 3 μm resolution. Probes features were extracted using the Feature Extraction Software v. 10.7.3.1 with GE_1_Sep09 protocol (Agilent Technologies). Intra-array normalizations were directly performed by the Feature Extraction Software. The raw data are available in the GEO database (GSE98328). Quantile inter-arrays normalization was performed using the Expander software (Sharan et al., 2003). Data were Log_2_ transformed and not available (NA) value was assigned if Found and/or Well Above Background flags were not positive. Expression values of probes with the same sequence were mediated. In myofiber microarray experiments, 10 myofibers per each of 6 myofiber types (classified by MyHC protein expression, total: 60 microarrays) were analyzed. Only probes with at least 8 of 10 available values in at least one myofiber type were taken into consideration for the further analyses. In cells microarray experiments, 6 independent replicates for miRNA mimic transfected cells, and 4 independent replicates for cells transfected with a scrambled sequence (used as control) were analyzed. Only probes with at least 75% of available values in at least one condition were taken into consideration for the further analyses.

#### Statistical analyses of microarray data

Microarray data were analyzed using the MultiExperiment Viewer (MeV, Ver. 4.8), a tool of TM4 Microarray Software Suite (Saeed et al., 2003). To identify the DE probes among myofibers, we performed one-way analysis of variance (ANOVA). Each sample was assigned at one of the 6 groups identified by the MyHC protein classification. Probes with an adjusted Bonferroni P-value (based on 1,000 permutations) lower than 0.01 were considered as DE. In addition, to improve the number of genes differentially expressed, probes not recognized by the statistical test and exclusively expressed in 1 or 2 clusters were manually added to the list. Samples (myofibers) were hierarchically clustered using the Pearson’s correlation distance and average linkage method. DE probes were grouped in 8 different clusters by K-means clustering, on the basis of their expression. Intensity values of the probes of the same cluster and of the same transcriptional class were mediate and differences among classes were considered statistically and biologically significant if the adjusted P-value generated by ANOVA was < 0.01 (calculated with GraphPad Prism software) and the difference of means > 1.

To identify the DE probes in transfected cells, we performed Significance Analysis of Microarrays (SAM), a non-parametric statistical test based on a permutation approach specifically implemented for microarray data, imposing a threshold level of False Discovery Rate of 0.05% (Tusher et al., 2001).

#### Gene ontology

DE genes were categorized in gene ontology (GO) classes using the Functional Annotation Clustering method, a bioinformatic tool of the DAVID database (Huang da et al., 2009). Only GO terms with an enrichment score greater than 1.3 and with P-value less than 0.05 were considered.

#### Analysis of discriminant genes

The list of the best marker genes among transcriptional myofiber classes was obtained applying Prediction Analysis of Microarray (PAM) algorithm (Tibshirani et al., 2002). PAM utilizes the shrinkage nearest centroid method that does automatic gene selection. Missing values are imputed using a K-nearest neighbor average in gene space (K = 10).

### mRNA q-RT-PCR

#### Primer design

qRT-PCR primers (**Table S4**) were designed by the primer design program Primer3Plus. All primer pairs span intron-exon boundaries. The specificity of the each amplicon was assessed by dissociation curve analysis.

#### Reverse transcription

First-strand cDNA was synthetized following SuperScript III (Thermo Fisher Scientific) protocol in a total volume of 30 μl. Oligo-dT was used for reverse transcription for all the samples.

#### q-RT-PCR reaction

SYBR Green technology was applied for all the assays with ABI 7500 Standard q-RT-PCR System (Thermo Fisher Scientific). The total reaction volume was 10 μl, including 5 μl 2X GoTaq qPCR Master Mix (Promega), 0.6 μl of 10 μM left primer, 0.6 μl of 10 μM right primer, 1 μl cDNA template, and 2.8 μl of water. Each assay was performed in triplicate. Negative controls without template were added each time. The PCR program started with 2 min. at 95°C, followed by 40 cycles of two temperature steps (95°C, 15 sec.; 60°C, 1 min.) and ended with 15 sec. at 95°C, 1 min. at 60°C, 15 sec. at 95°C and, 15 sec. at 60°C. The last steps of PCR are performed to acquire the dissociation curve, validating the specificity of the PCR products. The threshold cycle (Ct) is defined as the fractional cycle number at which the fluorescence exceeds the fixed threshold of 0.2. Three replicates were performed for each reaction. To evaluate differences in mRNA expression, a relative quantification method was chosen where the expression of the mRNA target is standardized by the Txn1 endogenous mRNA, whose level remains essentially constant among different samples (Chemello et al., 2011).

### miRNA qRT-PCR

#### Reverse transcription

Volume of miRNA population purified from a single myofiber was reduced by speed vacuum concentrator to 6 μl and split in two portions. First-strand cDNA was synthetized using the Megaplex RT Primers Rodents Pools A and B (Thermo Fisher Scientific) and the TaqMan MicroRNA Reverse Transcription Kit (Thermo Fisher Scientific) following the corresponding protocol.

#### Preamplification reaction

As miRNA amount purified from a single myofiber is very low, specific cDNA targets were amplified to increase the quantity of desired cDNA for expression analysis. Preamplification was performed using TaqMan PreAmp Master Mix Kit (Thermo Fisher Scientific) and Megaplex PreAmp Primers Rodents Pools A and B (Thermo Fisher Scientific) in accordance with the manufacturer’s instructions. For each transcriptional signature, pools of three cDNAs from isolated myofibers were prepared for q-RT-PCR experiments.

#### qRT-PCR

qRT-PCR reactions were performed using TaqMan Universal Master Mix II, no UNG (Thermo Fisher Scientific) and a different TaqMan Small RNA Assay (Thermo Fisher Scientific) for each miRNA analyzed. Three replicates were performed for each reaction. Small nuclear RNA U6 was used as endogenous control, and negative controls without template were added in each plate. Reactions were run in ABI 7500 Standard q-RT-PCR System (Thermo Fisher Scientific) with the following program: 95°C, 10 min; 35 cycles of 95°C 15 sec. and 60°C 1 min. Relative quantification results were obtained using the ExpressionSuite Software (Ver. 1.0, Thermo Fisher Scientific). Pearson’s correlation with sequencing data was performed with Excel.

### NGS of miRNAs purified from single myofibers

#### Preparation of libraries

miRNAs libraries preparation partially follows previously set up method for miRNA labeling (Biscontin et al., 2010). Volume of miRNA population purified from a single myofibers was reduced by speed vacuum concentrator to 6.5 μl. miRNAs were polyadenylated at 3’ end using the Poly(A) Tailing Kit (Thermo Fisher Scientific), scaling the volumes of reagents in order to performed the reaction in 10 μl. Polyadenylated miRNA molecules were precipitated with NaOAc 3 M pH 5.5 (1/10 volume) and absolute ethanol (4 volumes) overnight at −20°C and resuspended in 3.2 μl of water. First-strand cDNA was synthetized using SuperScript II (Thermo Fisher Scientific) in a final volume of 10 μl, including 10 pmol of oligo-dT-Ion P1 Adapter primer (5’-CCTCTCTATGGGCAGTCGGTGATCCTCAGC[dT]20VN-3’) and 50 pmol of SMART primer (5’-CACACACAATTAACCCTCACTAAAggg-3’). ssDNAs having a SMART anchor sequence at the 5’-end was exponentially amplified by 10 PCR cycles performed with Platinum Taq DNA Polymerase High Fidelity (Thermo Fisher Scientific) using as right primer P1 and left primer an A Adapter primer (5’-CCATCTCATCCCTGCGTGTCTCCGACTCAG-3’) plus a barcode sequence specific for each myofiber type (tS: AAGAG; tI: TACCA; tF: CAGAA) plus SMART primer. A second step of PCR amplification was performed for 10 cycles using as primers A and P1. After each PCR reaction, products were purified twice through GenElute PCR Clean-up columns (Sigma-Aldrich) to eliminate primer-dimers. Library size selection was performed using E-Gel SizeSelect Gels (Thermo Fisher Scientific), according to manufacturer protocol, and recovering dsDNA longer than 120 nucleotides and shorter than 160.

#### Sequencing

Different tagged libraries were pooled together and used for emulsion PCR. Samples were loaded on the Ion OneTouch System (Thermo Fisher Scientific) according to manufacturer specifications and template-positive beads were purified through the Ion OneTouch ES (Thermo Fisher Scientific). Enriched beads were used to load 3 Ion PGM Chips (Thermo Fisher Scientific) and sequencing runs were performed using Ion PGM Sequencer (Thermo Fisher Scientific) according to manufacturer’s protocol.

#### Sequencing data processing

Raw reads were trimmed and cleaned-up using an in-house developed program that: a) identifies the barcodes of the different myofiber types; b) removes adapters; c) keeps only the reads longer than 18 nucleotides. The expression of miRNAs was quantized using miRDeep software (Friedlander et al., 2008). The processed reads were mapped to the known mouse miRNA precursors from miRBase database (Ver. 19) using the mapper module of miRDeep with default values. Basically in this process equal reads are counted and collapse. Reads that mapped more than 5 times are automatically excluded. Quantize module was used to normalize read counts of mature miRNAs.

#### Analyses of miRNA families

Enriched miRNA families analysis was performed on miRNAs obtained from NGS of myofibers using miR Family TargetScan database (Ver. 6.2). Only families with a P-value (Benjamini-Hochberg adjusted) lower than 0.05 and at least 2 identified miRNAs were considered statistically significant.

### Construction of mRNA-miRNA network

miRNA targets were identified using miRSVR (August 2010 release) and TargetScan (Ver. 6.2) prediction algorithms. Pearson’s correlation between mRNA (from microarray) and miRNA (from sequencing) expression data was performed with Excel (Microsoft Office). Only interactions with a correlation coefficient < 0.3 were considered anti-correlated. Networks were generated and analyzed using Cytoscape software (Ver. 3.2, (Shannon et al., 2003).

### miRNA mimics over-expression

C2C12 myoblasts were cultured in Dulbecco’s Modified Eagle Medium with high glucose (Life Technologies) + 10% Fetal Bovine Serum (Life Technologies) in 10 cm dishes and split every 2 or 3 days before they reached 70% of confluence. Cells were transfected using Lipofectamine 2000 (Life Technologies) following the manufacturer’s instructions with 150 nM of mirVana miRNA mimics or scrambled sequence, purchased from Life Technologies. Transfection efficiency was evaluated with Ambion Silencer FAM-Labeled Negative Control (Life Technologies) and estimated of about 60% by fluorescence microscopy and flow cytometry. Each transfection experiment was independently repeated at least in triplicate. Results were evaluated after 72 hours of transfection.

Hindlimb mouse muscles were transfected using Invivofectamine 2.0 (Life Technologies). Briefly, miRNA mimics were diluted to 0.5 mg/ml using complexation buffer, mixed to an equal volume of Invivofectamine, and incubated at 50°C for 30 min. Immediately 80 μl were injected in hindlimb muscles. Injection was repeated after 3 days and muscles were collected for analyses after one week from the first injection. Each transfection experiment was independently repeated at least in triplicate.

### Imaging

#### Immunostaining

1×10^4^ C2C12 myoblasts were grown on 13 mm round coverslip in 25 mM glucose DMEM or FA oxidation medium (for the complete description see Seahorse analyses) and transfected as indicated. Following 72 hours of incubation, cells were fixed for 15 min. at room temperature with 4% ice-cold formaldehyde. Mitochondria of cells were stained with previously co-transfection with mito-RFP or with anti-TOM20. For anti-TOM20 staining, cells were permeabilized for 10 min. with 0.5% Triton X-100, blocked for 1 hour with 3% BSA in PBS, and then stained overnight at 4°C with a rabbit polyclonal anti-TOM20 (1:150, Santa Cruz). After washing with PBS, cells were incubated with Alexa Fluor 568 Goat Anti-Rabbit IgG (1:1000, Life Technologies) for 1 hour at room temperature. Coverslips were mounted on glass slides using ProLong Gold Antifade Reagent (Life Technologies). Where indicated, neutral lipids were stained before mounting with 15 min. incubation with BODIPY 493/503 (2 μg/ml, Life Technologies). Each transfection was replicated independently five times. Images were acquired with a 60X objective using a confocal spinning-disk microscope (Andromeda iMIC system; TILL Photonics). Z-stacks images of ten randomly chosen fields for each coverslip were acquired and stored for subsequent analysis. Images were processed using ImageJ software. Quantification of the fragmentation index *f* of mitochondrial networks, defined as the sum of relative fragment areas that individually constitute less than 20% of the total mitochondrial area, was performed using MitoLoc software (Vowinckel et al., 2015). Statistical analysis was performed with GraphPad Prism software.

#### Muscle cryosection staining

Succinate dehydrogenase stain was performed incubating fresh muscle cryosections derived from *gastrocnemius* for 30 min. as described in (Blanco et al., 1988). Periodic acid-schiff (PAS) staining was performed on cryosections of *tibialis anterior* following the instructions of PAS staining system (Sigma-Aldrich). Images were acquired with a 10X objective using DMR Leica microscope. Bodipy staining was performed incubating cryosections of *tibialis anterior* with BODIPY 493/503 (2 μg/ml, Life Technologies) for 20 min. Images were acquired with a 60X objective using Zeiss LM700 Confocal. Images were processed using ImageJ software. Statistical analysis was performed with GraphPad Prism software.

#### Electron microscopy

Samples were fixed with 2.5% glutaraldehyde in 0.1 M sodium cacodylate buffer pH

7.4 for 1 hour at 4°C, post-fixed with 1% osmium tetroxide and 1% in 0.1 M sodium cacodylate buffer for 2 hours at 4°C. After three water washes, samples were dehydrated in a graded ethanol series and embedded in an epoxy resin (Sigma-Aldrich). Ultrathin sections (60-70 nm) were obtained with an Ultrotome V (LKB) ultramicrotome, counterstained with uranyl acetate and lead citrate and viewed with a Tecnai G2 (FEI) transmission electron microscope operating at 100 kV. Images were captured with a Veleta (Olympus Soft Imaging System) digital camera.

### Measurement of oxygen consumption

Oxygen consumption rate (OCR) was measured with the XF24 Extracellular Flux Analyzer (Seahorse Bioscience). C2C12 transfected myoblasts were seeded in XF24 cell culture microplates at 6 x 10^3^ cells/well in 0.2 ml of DMEM containing 25mM glucose and incubated at 37°C in 5% CO_2_. Experiments were carried out on confluent monolayers after 72 hours. OCR in 25 mM glucose DMEM was initiated by replacing the growth medium with serum-free DMEM glucose. OCR driven by lipid oxidation was initiated by replacing the growth medium with substrate-limited medium (DMEM with 0.5 mM Glucose, 1 mM glutamine, 0.5 mM carnitine and 1% FBS) 9 hours before the experiment. The medium was replaced with the FA oxidation (FAO) medium (2.5 mM Glucose, 0.5 mM carnitine, 111 mM NaCl, 4.7 mM KCl, 1.25 mM CaCl_2_, 2 mM MgSO_4_, 1.2 mM NaH_2_PO_4_, and 5 mM HEPES) 45 min. prior the assay. OCR driven by lipid oxidation was determined by inhibition of the mitochondrial FA importer CPT-I using 40 μM etomoxir added 15 min. before the experiment. Just prior to starting the assay XF Palmitate-BSA FAO substrate (Seahorse Bioscience, Part #102720-100) was added to the wells. A titration with FCCP was performed for each transfection in order to determine the optimal FCCP concentration (i.e., the concentration that stimulates respiration maximally), which was found to be 0.4-0.6-1 μM for scramble, 0.4-0.6-1 μM for miR-27a, 0.4-0.6-1 μM for miR-142. After OCR baseline was established, a solution containing oligomycin (in 25mM glucose experiment), FCCP, rotenone or antimycin A were sequentially added to each well to reach final concentrations of 1 μg/ml oligomycin, FCCP as stated above, and 1 μM for rotenone and antimycin A. Data are expressed as pmol of O_2_ per min. per plated cells. At the end of each experiment the medium was removed from each well and the total protein content per well was quantified. miRNA transfections were independently replicates at least three times, and for each one five technical replicates were measured. Statistical analysis was performed with GraphPad Prism software.

### Lipidomic analysis

#### LC-MS Measurements

Lipids were extracted from transfected cells using a solution 2:1 chloroform:methanol 0.005% BHT. The chloroformic extracts were taken, dried using a rotary evaporator, and dissolved in 150 μl HPLC-grade methanol (Sigma-Aldrich). MS measurements were carried out in both positive and negative ion mode, using a Bruker Esquire-LC quadrupole Ion-Trap mass spectrometer equipped with an electrospray source (Bruker Optik GmbH, Ettlingen, Germany). The spectrometer was coupled to an HPLC (Hewlett–Packard Model 1100 Series) to chromatographically resolve the analytes prior to their mass detection. The chromatographic separation was obtained using a Zorbax Eclipse XDB-C18 column (150 x 4.6 mm I.D., particle size: 3.5 μm; Agilent Technologies). The mobile phase consisted of solvent A, 70:30 methanol:water with 12 mM ammonium acetate, and solvent B, 100% methanol with 12 mM ammonium acetate. The elution program started from 35% B, reached 100% B in 40 min, and kept this composition for 50 min., at the steady flow rate of 0.8 mL/min. For the mass detection, the instrumental parameters were the following: scan range 100–1200 m/z with a frequency of 13000 m/z s^−1^, nitrogen pressure of 35 psi, temperature of 300°C, and high voltage capillary of either 4000 V (positive ionization mode) or –4000 V (negative mode). The injection volume was 10 μl.

#### Data Analysis

The lipid quantitation was performed by integration of the extracted ion chromatograms (XIC), achieved through the proprietary software Bruker Daltonics esquireLC 4.5. Overall, the following lipid classes were detected: lyso-phosphatidylcholines (LysoPC), sphingomyelins (SM), phosphatidylcholines (PC), phosphatidylethanolamines (PE), phosphatidylinositols (PI), diglycerides (DAG), triglycerides (TAG), cholesteryl-esters (CE) and cholesterol. In order to compare the results across samples, the signal areas were normalized to the number of cells relative to each sample, which were therefore used as a reference for the quantitation. Experiments were independently repeat in quadruplicate. Statistical analysis was performed with GraphPad Prism software.

### Luciferase assay

Myoblasts were transfected with miRNA mimics and 100 pg/μl of pmirGLO Dual-Luciferase miRNA Target Expression Vector (Promega) containing the target sequence or a control sequence. Assays were performed using the Dual-Luciferase Reporter Assay (Promega), measuring firefly and renilla luciferase activities with Turner Designs TD-20/20 Luminometer (DLReady). miRNA transfections were independently replicates at least three times. Statistical analysis was performed with GraphPad Prism software.

### Glycogen assay

Glycogen content in transfected cells was measured using the Glycogen assay colorimetric kit (Abcam) following the manufacturer’s instructions. Results derive from the sum of six independent experiments. Statistical analysis was performed with GraphPad Prism software.

### Flow cytometry

Transfected C2C12 myoblasts were permeabilized and stained with BODIPY 493/503 (2 μg/ml, Life Technologies) for 15 min. After two PBS washes, cells were transferred in FACS conical tubes and flow cytometry was performed using a BD FACSCalibur platform (Becton Dickinson). Percentage of positive cells was calculated using mock-transfected C2C12 as a negative control. Results derive from the sum of six independent experiments. Statistical analysis was performed with GraphPad Prism software.

## SUPPLEMENTAL INFORMATION

**Figure S1:**
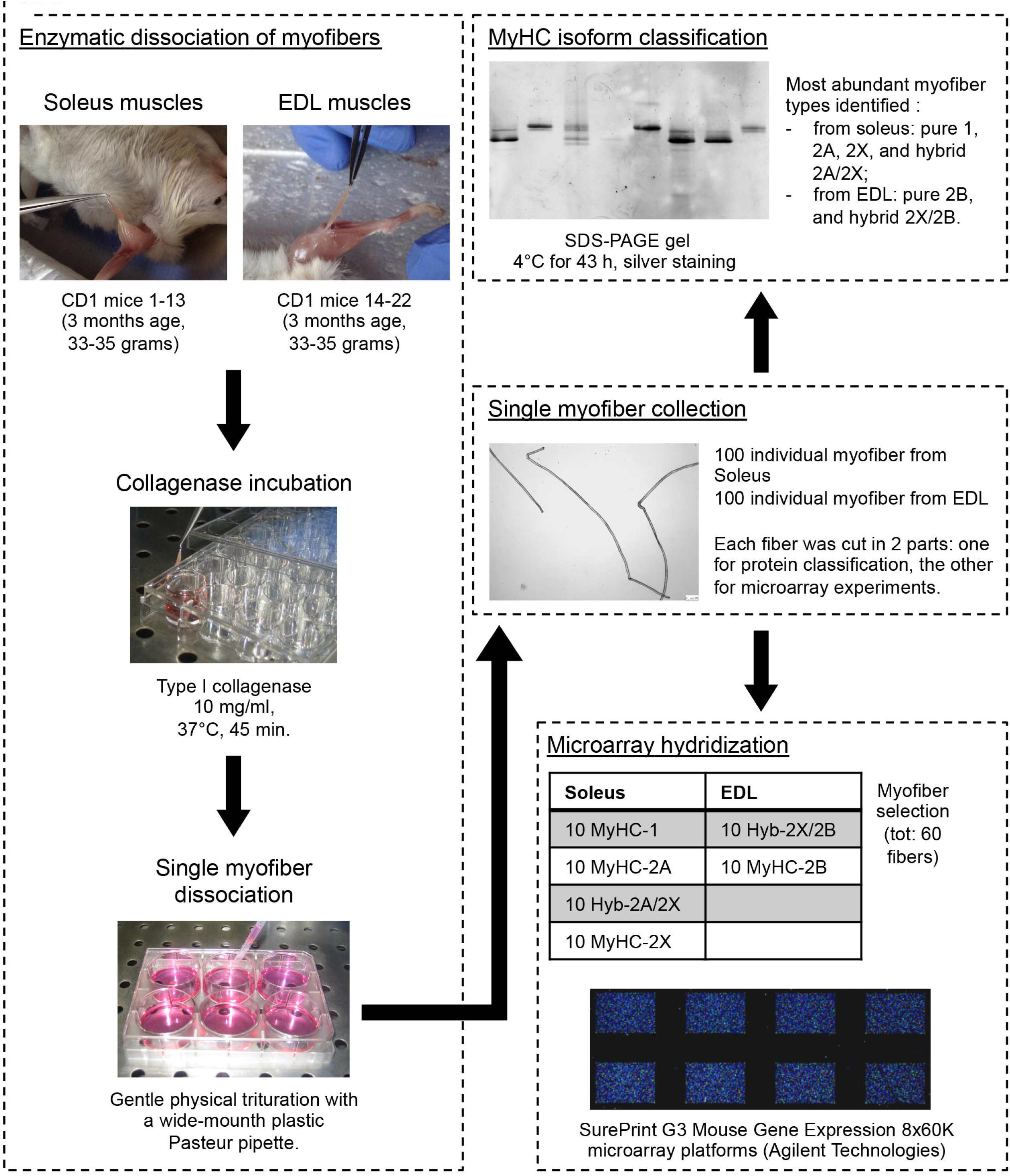
Single myofiber isolation and characterization. Myofibers were dissociated from *soleus* and EDL muscles of wild-type CD1 male mice at 3 months of age (Charles River, weight: 33 – 35 g). *Soleus* muscles were collected from mice numbered from 1 to 13 and EDL from mice numbered from 14 to 22. Muscles were separately moved in 1 ml high-glucose DMEM containing type I collagenase. Incubation proceeded for 45 min. at 37°C. The collagenase-treated muscles were rinsed and single myofibers were liberated by gentle physical trituration with a wide-mouth plastic Pasteur pipette. Intact, non-contracted, and well-isolated myofibers were picked under stereomicroscope and washed. We collected 100 myofibers from *soleus* muscles and the same number from EDL muscles. About one-third of each fiber was cut and placed in Laemmli buffer for fiber typing while the remaining part was dissolved in TRIzol Reagent for RNA purification. Fiber typing was performed using SDS-PAGE to identify MyHC protein isoforms. After silver staining, bands of MyHC appeared separated and were identified according to their migration rates. As expected, myofibers from *soleus* muscles were mainly pure 1, 2A, 2X, or hybrid 2A/2X, whereas myofibers from EDL muscles were mainly pure 2B or hybrid 2X/2B. For RNA analysis, we selected 60 myofibers, 10 for each most common myofiber types (biological replicates). RNA was amplified and labeled for microarray hybridization.

**Figure S2:**
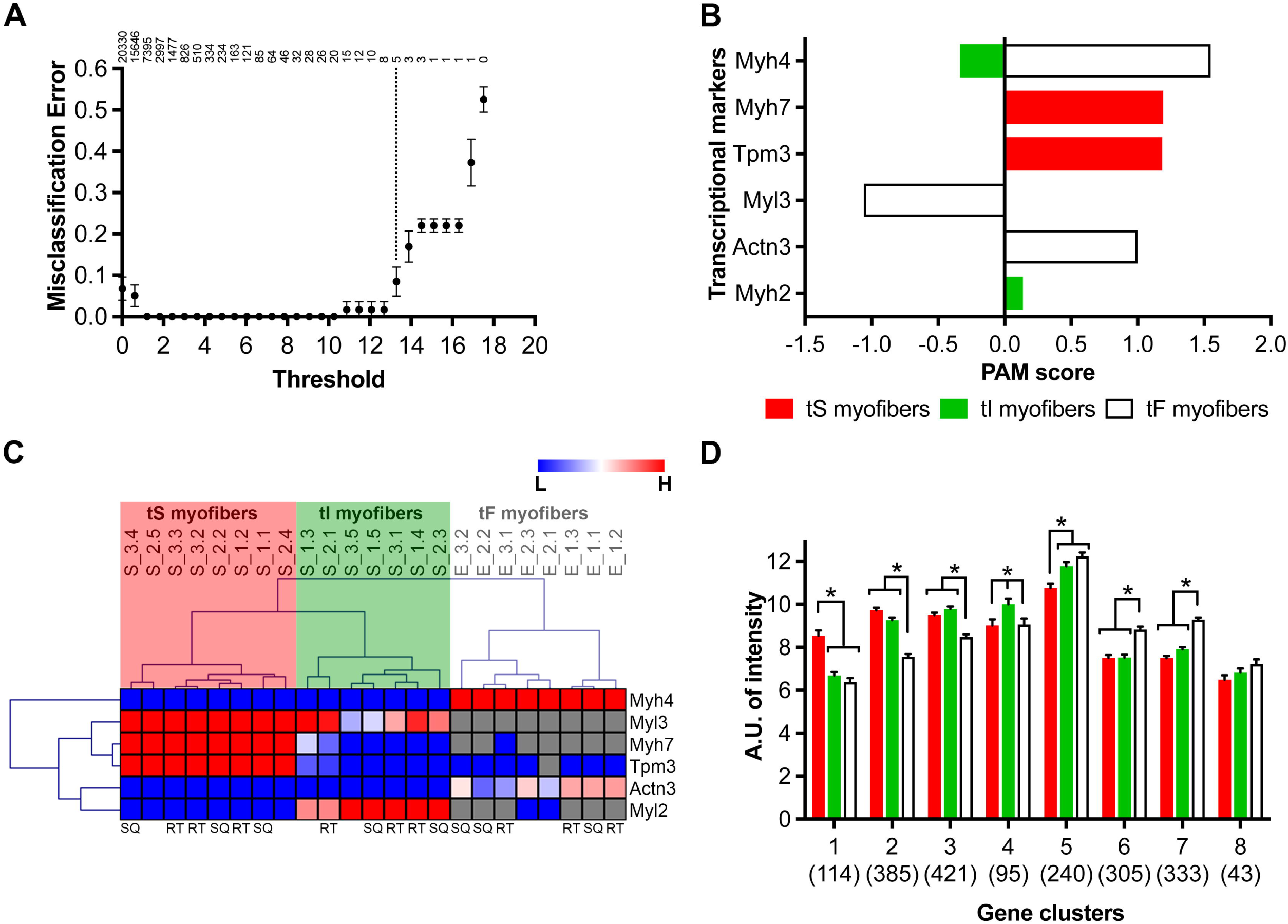
Characteristics of myofiber transcriptional signatures. (A) PAM test was used for showing the relationship between misclassification errors and number of genes used for building a classification model. The plot obtained by running PAM shows that, using a threshold value of 13.3, the expression of 5 genes (dotted line, Myh4, Myh7, Tpm3, Myl3, Actn3) can discriminate among fiber types with complete accuracy. The cross-validation error rate increases when too few or too many genes are involved in this prediction model. Number of genes is indicated at the top. (B) The best genetic markers for the classification of mouse myofibers at transcriptional level, with the assignment of specific scores for *soleus* or EDL muscles. The higher is the absolute value, the stronger is the classification power of the marker. For each marker, PAM score could be positive (highly expressed in a specific signature) or negative (low or not expressed in a specific signature). Myh2 was arbitrary added to the 5 genes identified by PAM analysis as the best marker of tI myofibers. (C) Expression levels of these genes were measured by qRT-PCR in 15 novel and unclassified myofibers isolated from *soleus* and 8 from EDL from 3 different mice (S = *soleus*, E = EDL; the first number after the underscore indicates the different mice; the second number, after the dot, indicates the different myofiber). Heat-map represents the normalized expression level against the reference gene Txn1: blue color means low expression, red color high expression, and grey color that the gene expression value was below the threshold of detection. Myofibers and genes were hierarchically clustered using the Euclidean distance and complete linkage methods. This analysis is sufficient to properly classify a myofiber in one of the three transcriptional signatures. At least two myofibers per transcriptional signature were used for miRNAs sequencing (SQ), and three myofibers for miRNAs qRT-PCR validation (RT). (D) K-means clusterization grouped the 1,936 probes with similar expression patterns among the three transcriptional myofiber signatures in 8 different clusters. Expression values (arbitrary units, A.U.) of each probe were separately mediated for the different transcriptional signatures and plotted in the bar graph. The numbers of probes of each cluster are indicated in bold, error bars are SEM. Except for cluster 8, each probe cluster shows a preferential expression in one or two myofiber types (*: adjusted P-value < 0.01, and difference of means > 1): probes of cluster 1 are more expressed in tS myofibers, clusters 2 and 3 in tS and tI myofibers, cluster 4 in tI myofibers, cluster 5 in tI and tF myofibers, and clusters 6 and 7 in tF myofibers (additional information in Dataset S4).

**Figure S3:**
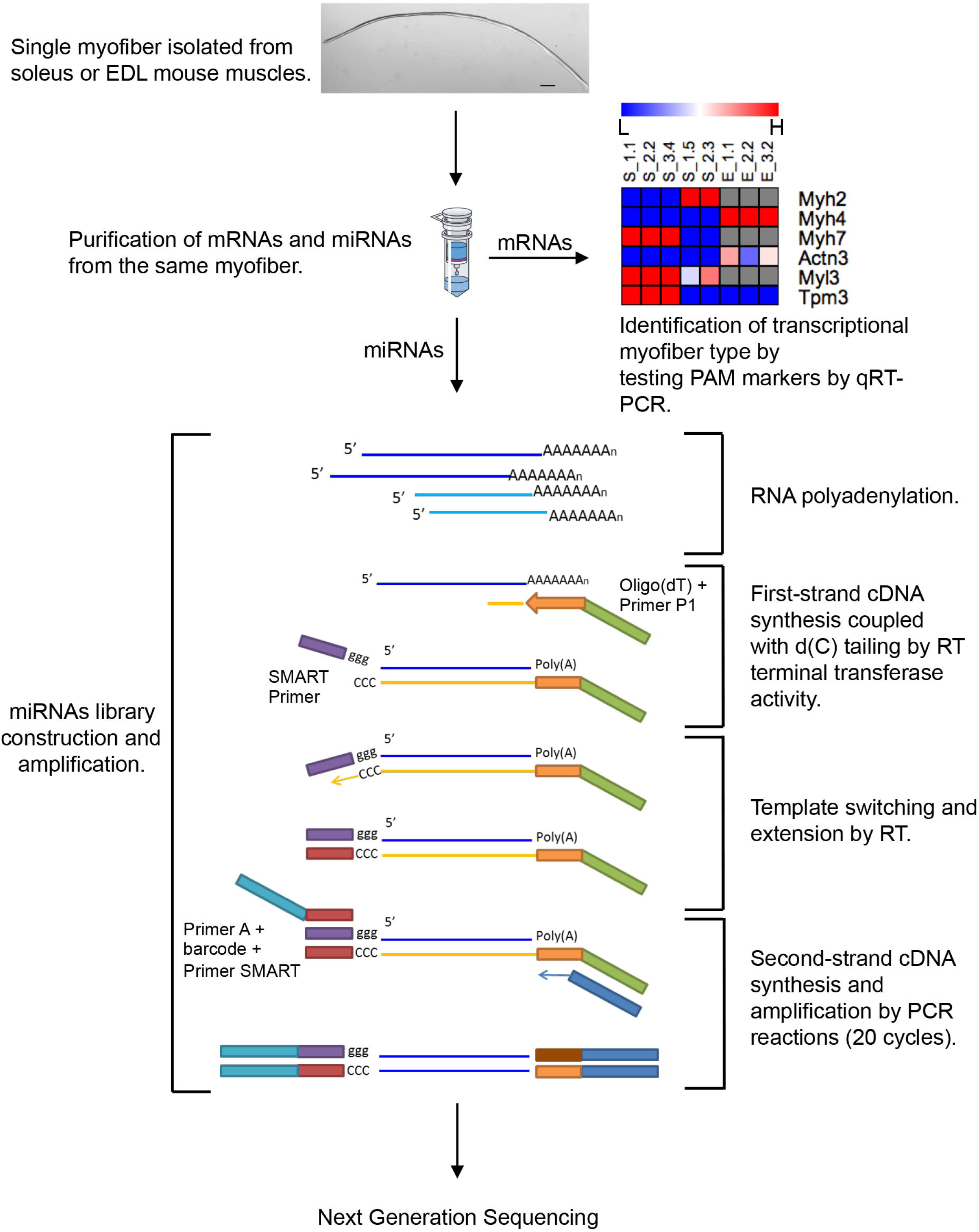
Workflow for the construction of miRNA libraries from single myofibers. From the same isolated myofiber (the picture shows an EDL myofiber, scale bar 250 μm), mRNA and miRNA populations were separately purified. mRNA was used for classifying myofibers according to the proposed transcriptional classification (tS, tI, or tF) by qRT-PCR. miRNA libraries were constructed and amplified using the Switching Mechanism at 5’ End of RNA Template (SMART) technology. Briefly, miRNAs were polyadenylated at 3’-end and then retrotranscribed with a modified anchored oligo(dT) primer. This presents on the 5’-end the P1 primer used for Ion Torrent emulsion PCR. Retrotranscription was performed using an enzyme that maintains terminal transferase activity allowing the association of SMART primer to the 3’-end of the cDNA. Second-strand cDNA and full-length miRNA libraries amplification were performed through PCR reaction. Primer P1 was used as reverse primer while as forward primer we used a sequence composed by (5’-3’ direction): the primer A for sequencing, a barcode sequence, and the SMART primer. PCR amplification was stopped after 20 cycles in order to achieve a good level of amplification without reaching the plateau phase. This approach allows a strand specific sequencing.

**Figure S4:**
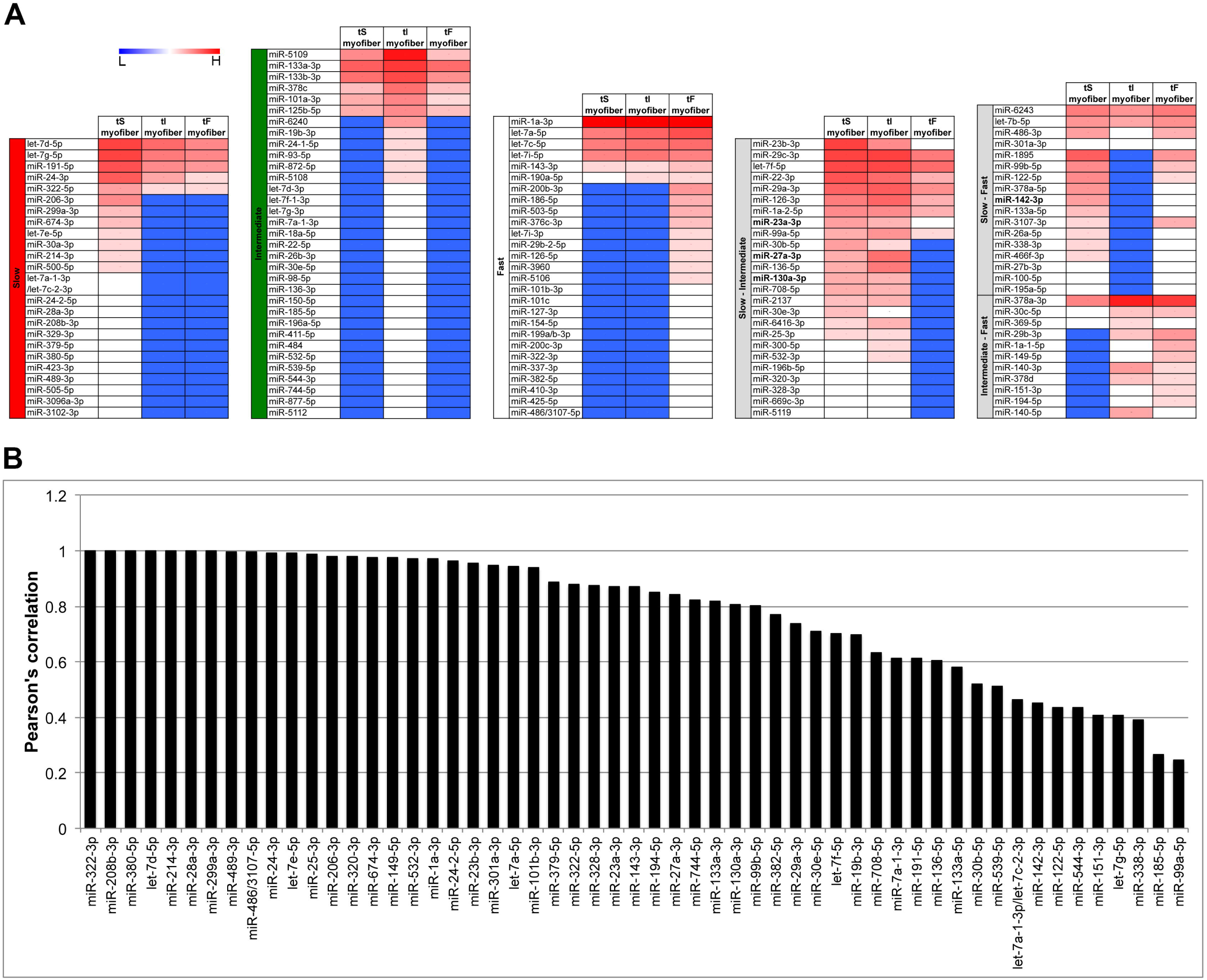
miRNome of different myofiber transcriptional types. (A) miRNAs identified in myofibers were grouped accordingly to their preferential expression by cluster analysis. Expression data are Log_2_ of sequencing results normalized as counts of reads per million (+1): blue is for poorly expressed (low, L), red for highly expressed (high, H). (B) The expression levels of 55 miRNAs among the three transcriptional myofiber types were measured by qRT-PCR and compared to NGS measurements. An averaged Pearson’s correlation of 0.78 was obtained between the two technologies, corroborating the reliability of NGS results.

**Figure S5:**
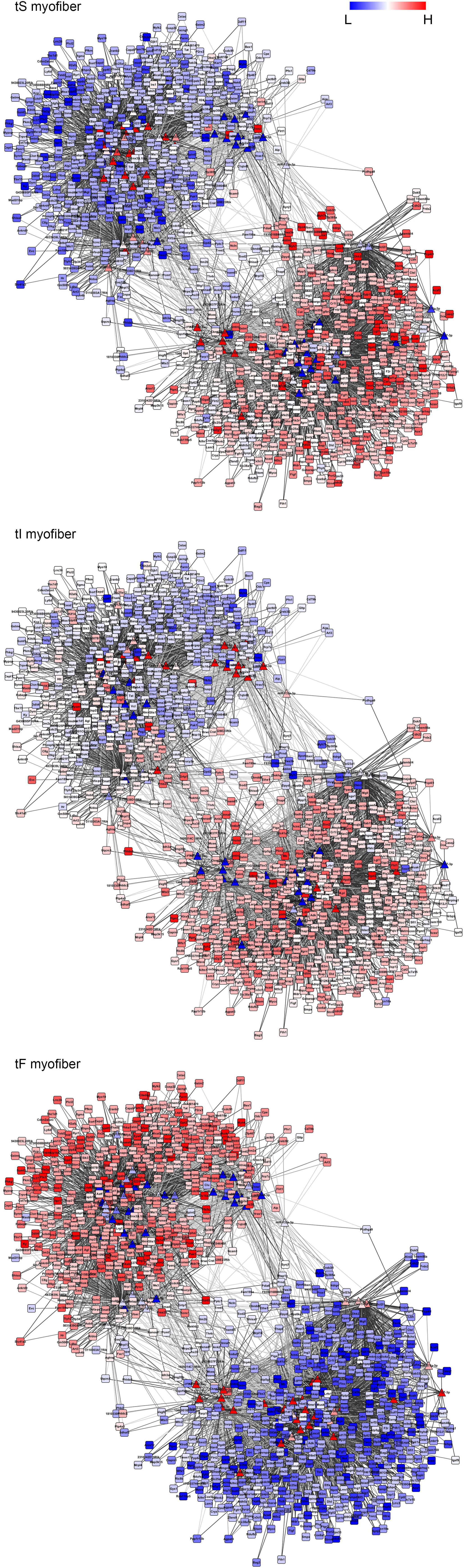
Post-transcriptional regulatory networks of myofibers. Data obtained from microarray (mRNAs) and NGS (miRNAs) experiments of single isolated myofibers were integrated to generate the post-transcriptional regulatory networks of tS, tI, and tF myofibers. miRNA targets were identified by TargetScan and/or mirSVR prediction algorithms. Only interactions with a correlation coefficient < −0.3 were considered. Networks are composed by 1,130 nodes including 79 miRNAs (triangles) and 1,051 mRNAs (squares): blue is for poorly expressed (low, L), red for highly expressed (high, H). They are connected by 5,968 edges (grey to black lines, with color intensity increase paralleling the increase of the anti-correlation between miRNA and mRNA target).

**Figure S6:**
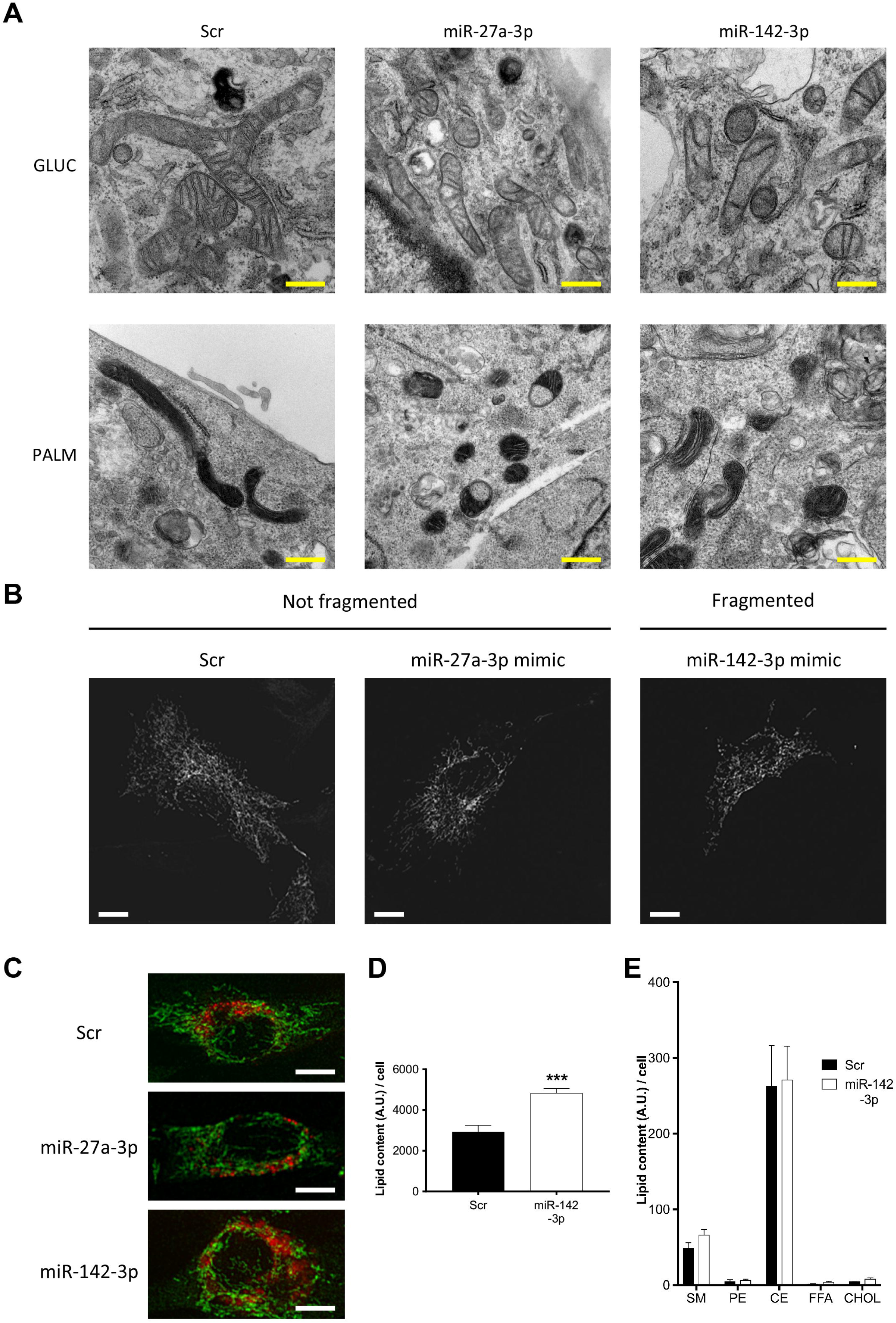
*In vitro* mitochondrial fragmentation induced by miR-27a-3p and −142-3p. (A) Electron microscopy of mitochondria of C2C12 myoblasts transfected with a scramble sequence as control (Scr), with miR-27a-3p, or with miR-142-3p in medium containing glucose (GLUC) or palmitate (PALM) as substrate. Scale bar is 500 nm. (B) Representative fluorescence microscopy images of transfected myoblasts in palmitate medium with different levels of mitochondria fragmentation. Mitochondria were stained with mRFP and imaged by confocal microscopy. Images of each cell were captured at different focal depths and then processed by focal plane merging and convolution. Scale bar is 10 μm. (C) Representative fluorescence microscopy images of single myoblasts. Mitochondria were stained with anti-TOM20 (green, mitochondria) and BODIPY (red, lipids) and analyzed by confocal microscopy. Scale bar is 10 μm. (D) Global lipid content of myoblasts transfected with a control scramble sequence or with miR-142, and measured with liquid chromatography-mass spectrometry (LC-MS) technology. Lipid quantification was measured as arbitrary units (A.U.) of signal areas of chromatograms and normalized to the number of cells relative to each sample. Number of independent experiments per condition *n* = 4. Error bars are SEM and adjusted P-values of ANOVA statistical test was indicated as follow: *** < 0.001, and it is referred in comparison to the control. (E) The figure shows the LC-MS quantification of classes of lipids (SM: sphingomyelin, PE: phosphatidylethanolamine, CE: cholesteryl ester, FFA: free fatty acid, CHOL: cholesterol) that show similar concentrations in myoblasts over-expressing miR-142-3p in comparison to control. Number of independent experiments per condition *n* = 4. Error bars are SEM; P-values of t-test are > 0.05.

**Dataset S1: MyHC protein and mRNA content of the 60 myofibers sampled for microarray experiments.**

Myofibers for microarray experiments were dissociated from *soleus* and EDL muscles taken from different mice. Fiber replicates of the same MyHC type were numbered from 1 to 10 and marked with muscle of origin (S for *soleus*, and E for EDL) and number of the animal. About one-third of each fiber was solubilized in Laemli buffer and used for MyHC protein isoform identification by SDS-PAGE. The relative proportions of MyHC isoforms were determined by the measurement of the brightness-area product (i.e., the product of the area of the band by the average brightness, with the subtraction of local background after black-white inversion). The remaining part of the fiber was immersed in TRIzol Reagent for RNA purification and amplification. Expression levels of Myh genes were measured by microarray experiments. Values correspond to Log_2_ spot fluorescence intensity after data normalization (see Materials and Methods). NA indicates that fluorescence of that specific microarray probe is above the detection limit.

**Dataset S2: Microarray probes differentially expressed among myofibers.**

Expression profiles were obtained using the Agilent microarray platform and technology. Values of arbitrary units of intensity were Log_2_ transformed and not available (NA) value was assigned if Found and/or Well Above Background Agilent Feature Extraction flags were not positive. Totally, 1,936 DE Agilent probes were identified: a) performing ANOVA one-way analysis, 1,730 probes with adjusted Bonferroni P-value (based on 1,000 permutations) lower than 0.01 were considered DE. The 6 groups considered in the ANOVA analysis corresponded to the 6 myofiber types identified by MyHC protein isoform classification (10 replicates for each group: MyHC-1, −2A, −2X, −2B, and Hyb-2A/2X, −2X/2B); b) manually selected probes not recognized by the statistical test and exclusively expressed in one or two transcriptional signatures. A probe is considered expressed in a particular transcriptional signature if the NA number is lesser than 40% and not expressed if the NA number is more than 75%. 206 probes satisfied these requirements (sheet “DE probes manually recovered”).

**Dataset S3: PAM test.**

Prediction Analysis for Microarray (PAM) was applied to microarray data to identify a small set of transcriptional markers useful for a quick myofiber classification (tS, tI, and tF). Imposing a threshold of 9.0, the list of the top 29 PAM markers was generated. Each PAM-selected mRNA could be a positive marker if it is expressed only in a particular transcriptional signature (PAM score > 0, red color) or a negative marker if it is not expressed in a particular transcriptional signature (PAM score < 0, green color).

**Dataset S4: Clusters of differentially expressed probes.**

K-means clustering on genes grouped the 1,936 differentially expressed Agilent probes in 8 different clusters, on the basis of their expression. Probes of cluster 1 (n = 114) are more expressed in tS myofibers, probes of clusters 2 (n = 385) and 3 (n = 421) in tS and tI myofibers, probes of cluster 4 (n = 95) in tI myofibers, probes of cluster 5 (n = 240) in tI and tF myofibers, and probes of clusters 6 (n = 305) and 7 (n = 333) in tF myofibers. Probes of cluster 8 (n = 43) do not show preferential expression in any transcriptional signatures.

**Dataset S5: Gene Ontology of genes differentially expressed in the 3 transcriptional myofiber signatures.**

DAVID bioinformatic tool v6.7 (http://david.abcc.ncifcrf.gov/) was used for Gene Ontology (GO) analysis of the differentially expressed Agilent probes that share common expression patterns among the 3 transcriptional myofiber signatures (tS, tI, and tF). In total, 1,893 unique probes (clusters 1 to 7) were used for the analysis and 1,451 were associated with a DAVID ID. The global set of genes in the mouse genome was used as background. One representative GO term from each functional annotation cluster with an enrichment score cutoff > 1.3 and P-values < 0.05 was listed. Overlapping functional clusters were ignored. Gene names for each group were listed.

**Dataset S6: miRNA identification from isolated myofibers by NGS.**

miRNA populations purified from single isolated myofibers were amplified following SMART technology protocol and sequenced using Ion Torrent PGM Sequencer. Libraries from miRNAs extracted from different myofibers belonging to the 3 transcriptional signatures were univocally flagged with 3 distinct sequence barcodes, in order to analyze in the same sequencing chip more than one library. miRNA libraries of one tS and one tF myofiber were placed and sequenced in a single Ion 314 Chip, and miRNA libraries from other 2 tS, 2 tI, and 2 tF myofibers were located and sequenced in further one Ion 316 Chip and one Ion 318 Chip. Raw reads were trimmed and cleaned using a proprietary program that identifies the barcodes of the different myofibers, removes the adapters, and keeps only the reads longer than 18 nucleotides. This produced 1,464,908 processed reads for tS, 610,994 for tI, and 943,182 for tF myofibers. The processed reads were mapped to the known mouse miRNA precursors from miRBase (v. 19), the expression levels were quantized using mirDeep and indicated as counts per millions of reads (CPM). This analysis shows that 81 miRNAs are expressed in tS myofibers, 84 in tI myofibers, and 75 in tF myofibers, for a total of 137 miRNAs found in at least one myofiber transcriptional type.

**Dataset S7: miRNA families expressed in muscle fibers.**

Enriched analysis of miRNA families was performed on myofiber miRNA sequences produced by NGS, using the miR Family TargetScan database (Ver. 6.2). Only families with a P-value (Benjamini-Hochberg adjusted) lower than 0.05 and at least 2 identified miRNAs were considered statistically significant and listed. Families containing known myomiRs are marked in red color.

**Dataset S8: miRNA qRT-PCR of single myofibers.**

55 miRNAs identified by NGS were tested by qRT-PCR in the 3 different myofiber types. mRNA and miRNA populations were separately purified: mRNA fractions were used for myofiber transcriptional classification and then miRNAs of three myofibers per transcriptional type (tS, tI, and tF) were used for qRT-PCR experiments. U6 was used as endogenous reference to calculate ΔCT values.

**Dataset S9: miRNA-mRNA targets network.**

To identify miRNA putative targets, we integrated for the analysis two complementary algorithms: a) TargetScan (version 6.2, release June 2012), that predicts biological targets of miRNAs by searching for the presence of conserved sites that match the seed region; b) mirSVR (release August 2010), a machine learning method able to identify also non-canonical and non-conserved miRNA target sites. In addition, only interactions with a negative Pearson’s correlation coefficient (< −0.3) between miRNA and mRNA expressions among the three transcriptional signatures were considered. miRNA expression values were reported as Log_2_ of count per million + 1, while mRNA expression values were reported as the mean of Log_2_ of the arbitrary units of microarray intensities. Values of the different probes identifying the same transcript were mediated. Each interaction was predicted by one or both prediction algorithms (TS = TargetScan, SVR = mirSVR, TS-SVR = both). The sheet “Number of direct edges” reports the number of edges for each node.

**Dataset S10: Lipidomic analysis.**

Lipids were extracted from at least half million of myoblasts transfected with a scrambled sequence (Scr, as control), or with miR-142-3p, and measured with LC-MS. In order to compare the results across samples, the signal areas were normalized to the number of cells relative to each sample, which were therefore used as a reference for the quantitation (*n* = 4). 66 lipids were identified (carbons:double bounds), belonging to 10 different classes. Lyso-PC: lysophosphatidylcholine, SM: sphingomyelin, PC: phosphatidylcholine, PE: phosphatidylethanolamine, PI: phosphatidylinositol, DAG: diacylglycerol, TAG: triacylglycerol, CE: cholesteryl ester, FFA: free fatty acid, CHOL: cholesterol.

**Dataset S11: Gene Ontology categories down-regulated in myoblasts over-expressing miR-27a-3p or miR-142-3p.**

As miRNAs repress the expression of their biological targets, Gene Ontology (GO) enrichment analysis of down-regulated genes in myoblasts over-expressing miR-27a-3p or miR-142-3p was performed using the DAVID bioinformatic tool. A single representative GO term from each functional annotation cluster with an enrichment score > 1.3 (the higher, the better) and P-values < 0.05 (the lower, the better) was listed. Overlapping functional terms were ignored.

